# Engineering of ATP synthase for enhancement of proton-to-ATP ratio

**DOI:** 10.1101/2024.08.27.609901

**Authors:** Hiroshi Ueno, Kiyoto Yasuda, Norie Hamaguchi-Suzuki, Riku Marui, Naruhiko Adachi, Toshiya Senda, Takeshi Murata, Hiroyuki Noji

## Abstract

F_o_F_1_-ATP synthase (F_o_F_1_) interconverts the energy of the proton motive force (*pmf*) and that of ATP via mechanical rotation of the rotor complex. The H^+^/ATP ratio, one of the most crucial parameters in bioenergetics, varies among species due to the different number of H^+^-binding *c*-subunits, resulting in H^+^/ATP ratios ranging from 2.7 to 5. The present study attempted to enhance the H^+^/ATP ratio significantly by employing a novel approach that differs from that of nature. We engineered F_o_F_1_ to form multiple peripheral stalks, each bound to a proton-conducting *a*-subunit. Engineered F_o_F_1_ showed an H^+^/ATP ratio of 5.9, beyond the highest among naturally occurring F_o_F_1_s, enabling ATP synthesis at a low *pmf*, at which wild-type enzymes are unable to synthesize ATP. Structural analysis showed that the engineered F_o_F_1_ formed up to three peripheral stalks and the *a*-subunits. This study not only provides important insights into the H^+^-transport mechanism of F_o_F_1_ but also opens the possibility of engineering the foundation of cell bioenergetics.

## Main text

F_o_F_1_-ATP synthase (F_o_F_1_) is a ubiquitous enzyme found in the membranes of mitochondria, chloroplasts and bacteria. It synthesizes ATP from ADP and inorganic phosphate coupled with proton translocation across membranes along the proton motive force (*pmf*)^1–3^. F_o_F_1_ is a unique molecular motor complex composed of two rotary molecular motors: F_1_ and F_o_. Bacterial F_o_F_1_ shows the simplest subunit composition of *a*_1_*b*_2_*c_x_* (*x* varies among species) for F_o_ and α_3_β_3_γ_1_δ_1_ε_1_ for F_1_^4–6^ (Fig. 1a). F_o_ is a membrane-embedded molecular motor driven by *pmf*; when protons are translocated through the proton pathway in F_o_ along *pmf*, the multimeric rotor ring composed of *c*-subunits (termed *c*-ring) rotates against stator *ab*_2_ (Fig. 1b). F_1_ is the catalytic core domain of F_o_F_1_ for ATP synthesis and hydrolysis^7^. When isolated from F_o_, F_1_ acts as an ATP-driven molecular motor that rotates the γε rotor complex against the catalytic α_3_β_3_ stator ring coupled with ATP hydrolysis^8^. These two motors are coupled via two stalks: the peripheral stalk composed of the *b*_2_ dimer stalk and the δ subunit. The central rotor stalk is composed of a γε complex and a *c*-ring. Under ATP synthesis conditions where *pmf* is sufficient and the rotational torque of F_o_ exceeds that of F_1_, F_o_ rotates the γδ complex in F_1_ in the reverse direction of ATP hydrolysis, inducing the ATP synthesis reaction on the α_3_β_3_ ring ^9,10^. When the torque of F_1_ exceeds that of F_o_, F_1_ rotates the *c*-ring in F_o_ to enforce F_o_ to pump protons in the reverse direction, thereby generating *pmf*. Thus, F_o_F_1_ interconverts the *pmf* and chemical potential of ATP hydrolysis via mechanical rotation.

**Fig. 1.**
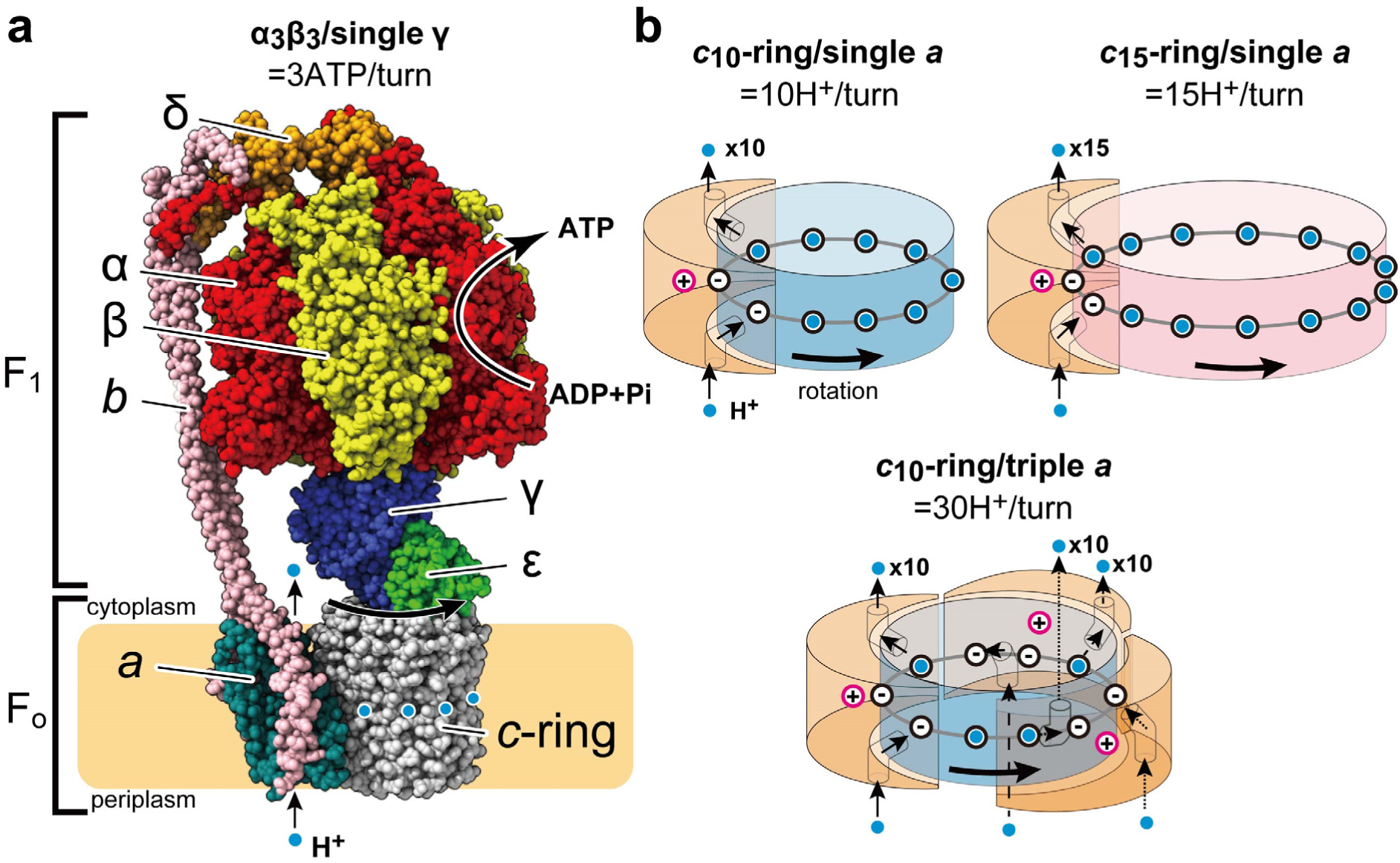
Structure of F_o_F_1_-ATP synthase and rotation mechanism of F_o_ with a different number of the *c*-/*a*-subunits. (a) Bacterial ATP synthase (thermophilic *Bacillus* PS3) consists of F_1_ (α_3_β_3_γδε) and F_o_ (*ab*_2_*c_10_*) motors. As F_1_ has three catalytic sites, three ATP molecules are synthesized per turn of the rotor subunits (γε*c_10_*) against the stator subunits (α_3_β_3_δ*ab*_2_) during ATP synthesis. (b) Models of proton translocation through F_o_ coupled with the rotation of the *c*-ring. The highly conserved arginine residues of the *a*-subunit and glutamate (or aspartate) residues of *c*-subunits are depicted with pink and black open circles, respectively. Protons are depicted as light blue circles. In the models of *c*_10_-ring/single *a*-subunit and *c*_15_-ring/single *a*-subunit, 10 and 15 protons, equal to the number of the *c*-subunits, are transferred in one turn, respectively. Conversely, in the *c*_10_-ring and triple *a*-subunit model, a total of 30 protons, equal to the number of the *c*-subunits multiplied by the number of the *a*-subunits, are transferred in one turn.

Considering the Gibbs free energy of this coupling reactions (Δ*G*’),

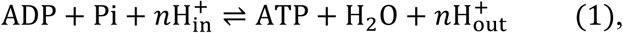

Δ*G*’ is given as:

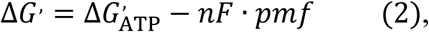

where Δ*G*^’^_ATP_ is the Gibbs free energy of ATP synthesis, *F* is Faraday’s constant, *n* is the H^+^/ATP ratio, which is defined as the number of protons translocated through F_o_ coupled with a single turnover of ATP synthesis on F_1_. Hence, the free energy of ATP synthesis reaction must satisfy the following condition:

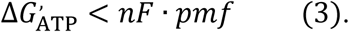

Thus, the H^+^/ATP ratio is the critical factor to determine the lower limit of *pmf* for ATP synthesis, considering that Δ*G*’ is not largely different among species. The H^+^/ATP ratio is principally defined from the ratio of the reaction stoichiometry of F_o_ per turn of the rotor complex to that of F_1_; and the ratio of H^+^/turn of F_o_ to ATP/turn of F_1_. All the F_1_s studied so far have three catalytic β subunits without exception, and couples three reactions of ATP hydrolysis/synthesis per turn, defining the ATP/turn as 3^10^. The reaction stoichiometry of F_o,_ H^+^/turn varies among species due to the variation in the number of *c*-subunits in the *c*-ring. According to the ‘half channel model’ supported by recent structural studies, the H^+^ pathway in F_o_ is formed by the *c*-ring and the *a*-subunit^11^ which has two half-channels, each exposed to the periplasmic or cytoplasmic side of the membrane^12,13^. During ATP synthesis, H^+^ in the periplasmic solution enters the half-channel exposed to the periplasmic space and the H^+^ is transferred to one of the *c*-subunits of the *c*-ring. After one revolution of the *c*-ring, H^+^ is released into the cytoplasmic solution via the opposite half-channel (Fig. 1b, Supplementary Fig. 1). Thus, the half-channel model assumes that the stoichiometry of H^+^/turn is determined by the number of the *c*-subunit.

The number of *c*-subunits of the *c*-ring in F-type ATP synthases ranges from 8 to 15 depending on the species^11^. When we assume the perfect energy coupling between F_1_ and F_o_, the H^+^/ATP ratio should vary between 2.7 and 5.0 among the species. Various groups have attempted to experimentally determine the H^+^/ATP ratio, from the biochemical measurement of thermodynamic equilibrium point where *pmf* and Δ*G*’_ATP_ is balanced. The *Bacillus* PS3 F_o_F_1_ with *c*_10_-ring has been reported to show good agreement with the structurally expected H^+^/ATP ratio of 3.3^14^. F_o_F_1_s from *E. coli* and yeast mitochondrial F_o_F_1_ which also have the *c*_10_-ring are reported to have slightly different H^+^/ATP ratios, 4.0 ± 0.3^15^ and 2.9 ± 0.2^16^, respectively. For spinach chloroplast F_o_F_1_ with the *c*_14_-ring, two independent studies reported smaller values, 4.0 ± 0.2^15^ and 3.9 ± 0.3^16^ than the expected value, 4.7. Thus, the experimentally determined H^+^/ATP ratios are close to, but not always the same as the structurally expected values, and vary in a narrow range, from 3 to 4.

The H^+^/ATP ratio of F_o_F_1_ is one of the most critical parameters in the bioenergetic system of cells, which defines the energy cost for ATP synthesis as well as the threshold for *pmf* for ATP synthesis (see Eq. 3). Since Δ*G*’ does not largely differ across organisms, F_o_F_1_ with a higher H^+^/ATP ratio is able to synthesize ATP even at lower *pmf*. In fact, alkaliphilic bacteria living in highly alkaline environments and photosynthetic organisms that grow under light-limiting conditions have a *c*-ring with a large number of *c*-subunits^17,18^. This is explained as an evolutionary adaptation to operate stable ATP synthesis reactions under low and/or unstable *pmf* conditions^19,20^. Thus, organisms may have optimized the H^+^/ATP ratio through evolution, by tuning the stoichiometry of the *c*-ring to meet their energetic requirements.

On the other hand, when reconsidering the half-channel mechanism, we can assume that the stoichiometry of the H^+^/turn of F_o_ is defined not only by the number of *c*-subunit but also by the number of *a*-subunit (Fig. 1b, bottom). In particular, the structures of F_o_ solved so far show that a large proportion of the *c*-ring is exposed to the lipid bilayer, indicating the possibility to accommodate another or two more *a-*subunits, although F_o_F_1_ with multiple *a*-subunits has not been found so far. In the present study, we sought this possibility to double or triple the H^+^/ATP ratio of F_o_F_1_, by multiplying the number of the *a*-subunits in F_o_.

## Results

### Design for multiple peripheral stalks

So far, all ATP synthases have a single copy of the *a*-subunit per F_o_F_1_ complex. Considering that the *a*-subunit is tightly bound to the membrane portion of the peripheral stalk, the key factor for multiplying the *a*-subunits should be the structural mechanism that limits the number of peripheral stalks. The peripheral stalk of the *b*_2_ dimer extends from the membrane to the upper surface of the α_3_β_3_ subcomplex of F_1_, binding to the δ subunit. The δ subunit binds onto the top of the α_3_β_3_ subcomplex, associating with the N-terminals of the three α subunits; one α subunit interacts with the C-terminal domain of the δ subunit, while the two interact with the N-terminal domain of the δ subunit as indicated with arrows in Fig. 2a. As the N-terminal domain of the δ subunit is located on the central concavity of the α_3_β_3_ ring, occupying the pseudo-rotational symmetry axis of the α_3_β_3_ ring, it is reasonable to assume that the N-terminal domain of the δ subunit breaks the pseudo-threefold symmetry, limiting the stoichiometry of the peripheral stalk to be one (Fig. 2a). We hypothesized that by removing the N-terminal domain of the δ subunit (Fig. 2b), it becomes possible to accommodate a truncated δ subunit on each α subunit (Fig. 2c). A possible concern regarding the truncation is that the truncated δ subunit (δ_ΔN_) does not form a stable complex with the α_3_β_3_ ring. Therefore, we designed the δ_ΔN_-α fusion construct of *Bacillus* PS3 F_o_F_1_, where the C-terminus of the δ subunit was genetically fused to the N-terminus of the α subunit. In addition, the inhibitory C-terminal domain of the ε subunit was removed for the enhancement of the activity^14,21^. In this study, *Bacillus* PS3 F_o_F_1_-ε_ΔC_ was used as a wild-type F_o_F_1_ for comparison (Supplementary Fig. 2).

**Fig. 2.**
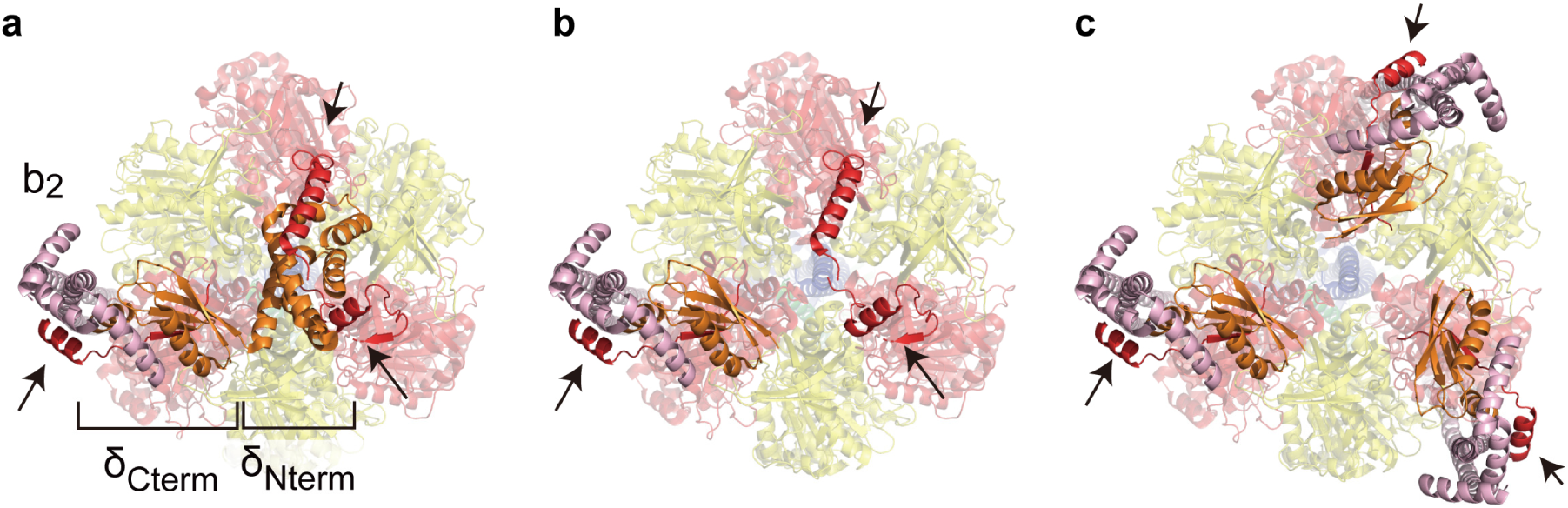
Design strategy to multiply peripheral stalks. Top views of *Bacillus* PS3 F_o_F_1_ from the cytoplasm (PDB ID: 6n2y). The α (red), β (yellow), γ (blue), δ subunits (orange), and the *b*_2_ stalk (pink) are shown in the cartoon. The non-transparent red parts indicated by the arrows represent the N-terminal region (residues 2–30) of the α subunit. (a) Asymmetric interactions between the δ and α subunits. The single δ subunit interacts with the three α subunits, occupying the central concavity of the α_3_β_3_ subcomplex. (b) Model structure after deletion of the N-terminal domain (residues 2–104) of the δ subunit. The central concavity is exposed, and two of three α subunits are unoccupied. (c) Model structure after each of the N-terminus of the three α subunits is fused to the C-terminal domain (residues 105–178) of the δ subunit.

### SDS-PAGE analysis of subunit stoichiometry

The δ_ΔN_-α fused F_o_F_1_ was purified following a previously reported procedure for the wild-type *Bacillus* PS3 F_o_F_1_^14^. To estimate the subunit stoichiometry, the δ_ΔN_-α fused F_o_F_1_ was analyzed by SDS-PAGE with the wild-type F_o_F_1_ for comparison (Fig. 3a, b). The δ_ΔN_-α fused F_o_F_1_ lacked δ and α, and the δ_ΔN_-α fusion appeared above the band position for the α subunit (Fig. 3a). The δ_ΔN_-α fused F_o_F_1_ retained the complete set of the subunits. Then, we estimated the subunit stoichiometry of the *b*- and *a*-subunits in the δ_ΔN_-α fused F_o_F_1_ by using the γ subunit as the internal reference in comparison with the wild-type (Fig. 3b). In the wild-type, the *b*-subunit showed comparable band intensity to the γ subunit while the *a*-subunit exhibited roughly half the intensity of the γ subunit. The δ_ΔN_-α fused F_o_F_1_ showed significantly higher signals for the *a*- and *b*-subunits relative to the γ subunit, indicating that the δ_ΔN_-α fused F_o_F_1_ increases the stoichiometry of the *a*- and *b*-subunits. For a more quantitative estimation, we drew calibration lines for the wild-type and mutant subunits and standardized the lines with the calibration lines of the γ subunit as the internal control (Fig. 3b). We then determined the stoichiometries of the *a*- and *b*-subunits of the δ_ΔN_-α fused F_o_F_1_ by comparison with the wild-type. The estimated stoichiometries of the *a*- and *b*-subunits were 2.4 and 2.0 times higher than those of the wild-type F_o_F_1_, respectively (see Fig. 3b legend). Thus, it was confirmed that the δ_ΔN_-α fused F_o_F_1_ has multiple, two on average, peripheral stalks and *a*-subunits.

**Fig. 3.**
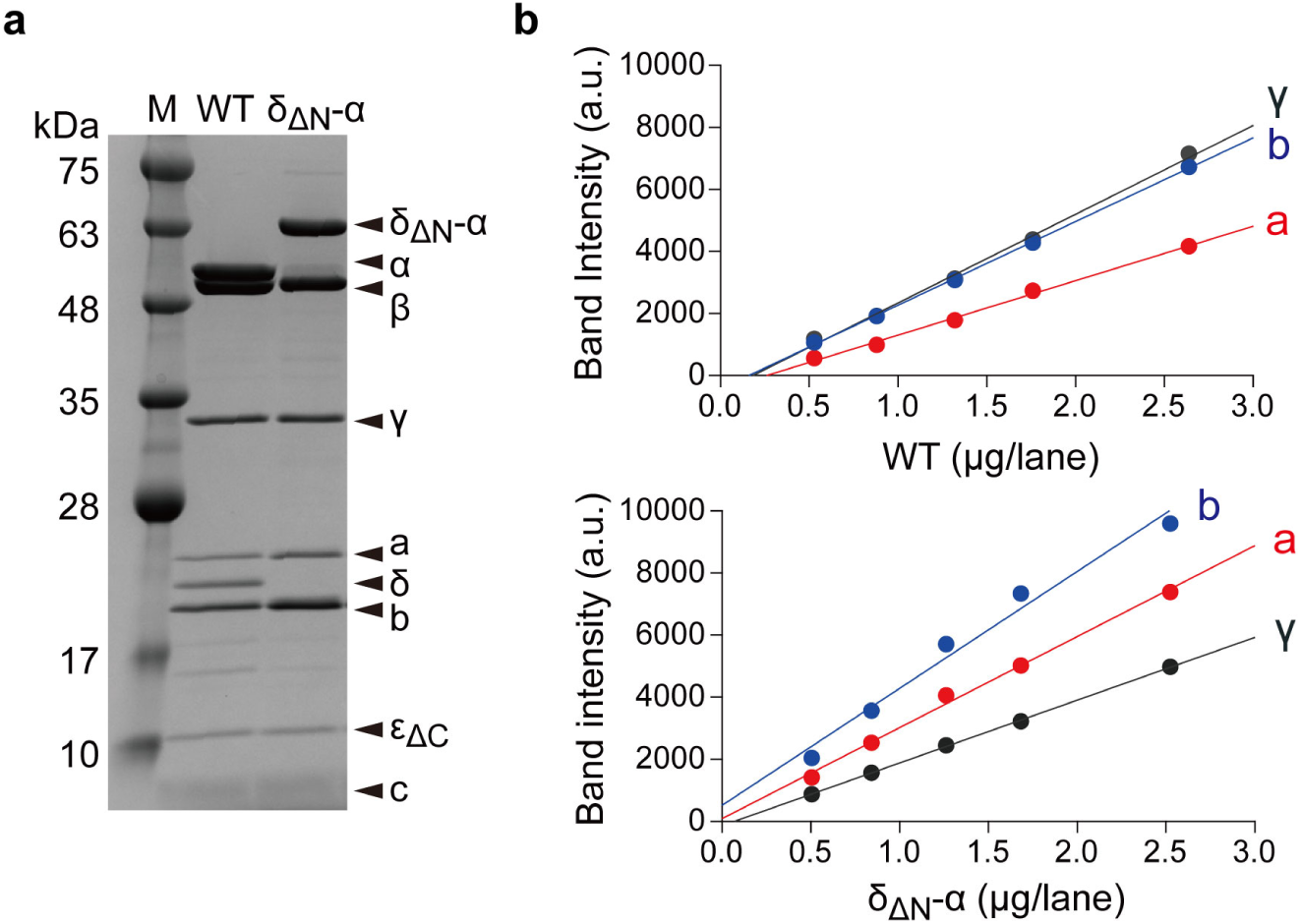
Subunit stoichiometry of δ_ΔN_-α fused F_o_F_1_. (a) SDS-PAGE analysis of the purified wild-type (WT) *Bacillus* PS3 F_o_F_1_ and δ_ΔN_-α fused F_o_F_1_. 3 μg of F_o_F_1_ was loaded in each lane. The molecular masses of the δ_ΔN_-α, α, β, γ, a, δ, b, ε_ΔC_, and *c*-subunits are 63, 55, 53, 32, 26, 20, 19, 9, and 7 kDa, respectively. (b) The band intensity vs total protein weight. The band intensity of each subunit was plotted against the total amount of F_o_F_1_ loaded for SDS-PAGE analysis. The plots were fitted with a linear function. The slopes for γ, *a-*, and *b*-subunits were determined to be 2861, 1757, and 2696 (a.u./µg) for WT F_o_F_1_, and 2021, 2929, and 3766 (a.u./µg) for δ_ΔN_-α fused F_o_F_1_, respectively. By normalizing the band intensity of the γ subunit of each F_o_F_1_, the stoichiometries of the *a*- and *b*-subunits of the δ_ΔN_-α fused F_o_F_1_ were estimated to be 2.4 (=(2929/2021)/(1757/2861)) and 2.0 (=(3766/2021)/(2696/2861)) times higher than those of the wild-type F_o_F_1_, respectively.

### Functional analysis of the H^+^/ATP ratio

We attempted to determine the H^+^/ATP ratio of the δ_ΔN_-α fused F_o_F_1_ from the biochemical measurement of thermodynamic equilibrium between *pmf* and Δ*G*’ as previously reported^14^. Firstly, we prepared the F_o_F_1_-reconstituted proteoliposomes (PLs) and incubated them in an acidic buffer. The PLs were injected into the base assay medium to initiate ATP synthesis. The ATP synthesis/hydrolysis activity was monitored with the luciferin/luciferase assay system under various *pmf* conditions, with a given reaction quotient, *Q* (= [ATP]⁄([ADP] · [Pi])). Fig. 4a shows the time courses of the assay, in which the initial rate was determined. Fig. 4b shows the reaction rates plotted against the *pmf* when *Q* = 2.5. The δ_ΔN_-α fused F_o_F_1_ was shown to catalyze the ATP synthesis reaction even at low *pmf*, at which wild-type F_o_F_1_ is unable to synthesize ATP. For a more quantitative analysis, the data points were fitted with an exponential function to determine the equilibrium *pmf* (*pmf*_eq_), where the torques of F_1_ and F_o_ are balanced and the net reaction rate is zero. At the condition of Fig. 4b where *Q* = 2.5, *pmf*_eq_ was determined to be 68 mV for the δ_ΔN_-α fused F_o_F_1_ and 133 mV for the wild-type. Thus, the minimum *pmf* for ATP synthesis was halved for the δ_ΔN_-α fused F_o_F_1_, suggesting that the functional H^+^/ATP ratio of the δ_ΔN_-α fused F_o_F_1_ is also doubled in agreement with SDS-PAGE analysis. For further confirmation, we determined the *pmf*_eq_ at various *Q* values (Supplementary Fig. 3). In all conditions, *pmf*_eq_ of δ_ΔN_-α fused F_o_F_1_ was almost half of that of the wild-type.

**Fig. 4.**
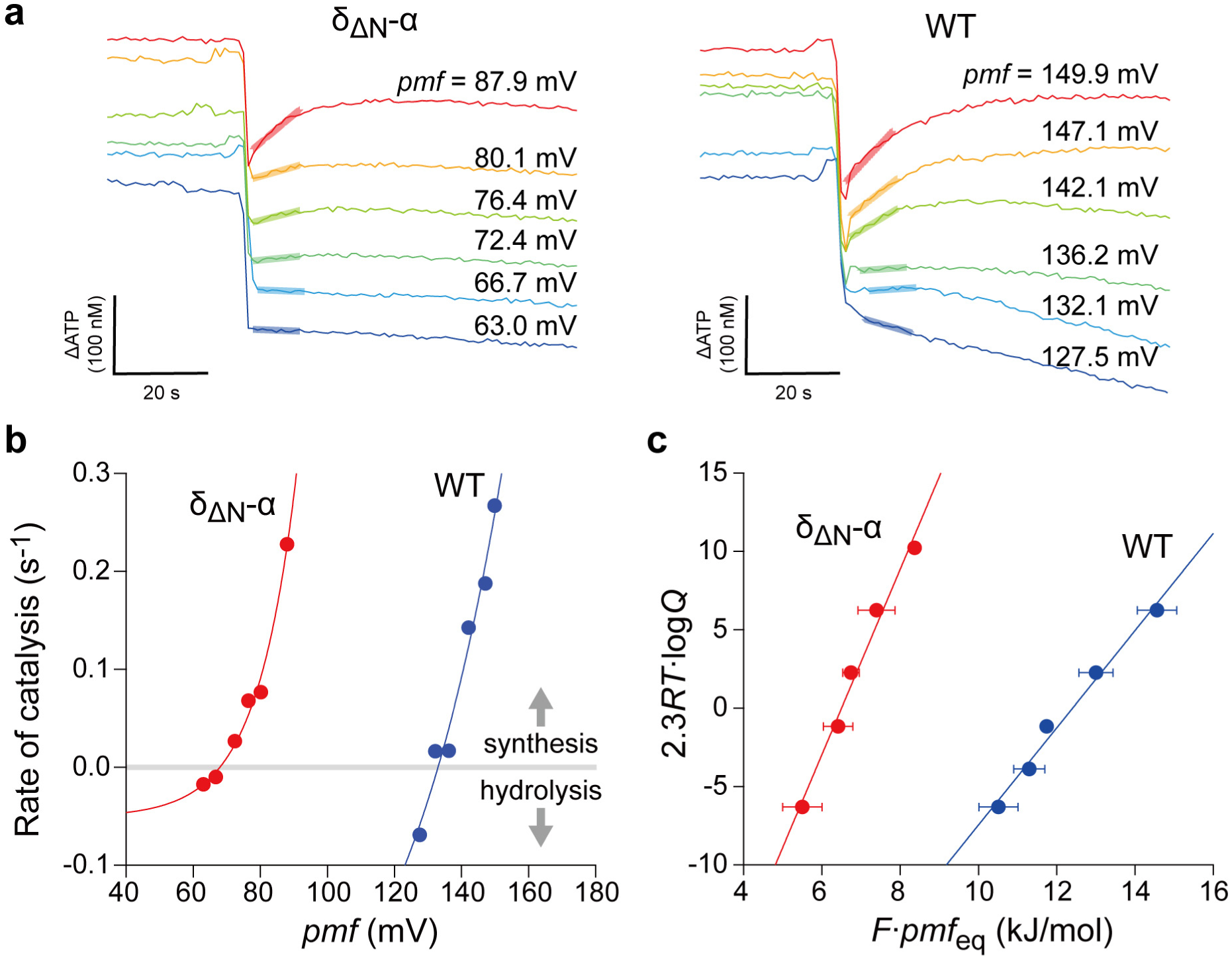
Functional analysis for the determination of the H^+^/ATP ratio. (a) Time courses of the ATP synthesis/hydrolysis activity of the reconstituted proteoliposomes (PLs) at different *pmf*. ATP synthesis reaction was measured using the luciferin/luciferase system. The reaction quotient, *Q* was 2.5; [ATP] = 500 nM, [ADP] = 20 µM, [Pi] = 10 mM. The rate of catalysis was determined from the initial slopes (bold lines). (b) The ATP synthesis/hydrolysis rates determined from (a) were plotted against *pmf*. The data points were fitted with an exponential function for the determination of the equilibrium *pmf, pmf*_eq_, as the interception of the *x* axis. (c) Determination of the H^+^/ATP ratio. Using all datasets shown in Supplementary Fig. 3, the mean and the SD of *pmf_eq_*values at each *Q* condition were determined, and 2.3*RT logQ* values were plotted against the corresponding *F* · *pmf_eq_* values according to Eq. (5). Each line represents a linear regression fit to the dataset obtained from each F_o_F_1_.

For the comprehensive analysis of the functional H^+^/ATP ratio, Eq. (2) was transformed as below,

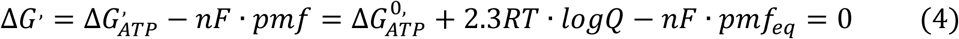

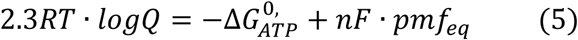

where 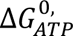 is the Gibbs free energy of ATP synthesis under the biochemical standard state, *R* and *T* are the gas constant and absolute temperature, respectively. Here, *pmf*_eq_ values were experimentally determined under defined *Q* conditions. The other values are constant. Therefore, when 2.3*RT logQ* is plotted against *F · pmf_eq_*, *n* (= the H^+^/ATP ratio) is determined as the slope of the data points. In addition, 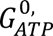 is determined as the interception of the y axis. As shown in Fig. 4c, the δ_ΔN_-α fused F_o_F_1_ exhibited a significantly steeper slope compared to the wild-type F_o_F_1_. From the linear fitting, the H^+^/ATP ratio was determined to be 5.9 ± 0.5 and 3.1 ± 0.2 (fitted value ± SE of the fit) for the δ_ΔN_-α fused F_o_F_1_ and wild-type, respectively. Although the H^+^/ATP ratio of the wild-type is slightly lower than the structurally expected value of 3.3 and the reported value (3.3 ± 0.1)^14^, δ_ΔN_-α fused F_o_F_1_ was shown to double the H^+^/ATP ratio, in close agreement with the subunit stoichiometry analysis from SDS-PAGE. This agreement suggests that the δ_ΔN_-α fused F_o_F_1_ has two functional *a*-subunits in the ensemble average. From the interception of the y axis, the 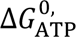 values are determined to be 38 ± 3 kJꞏmol^−1^ (fitted value ± SE of the fit) for both. This value shows a fine agreement with the reported values for *Bacillus* PS3 F_o_F_1_ (39 ± 1 kJꞏmol^−1^)^14^, *E. coli* (38 ± 3 kJꞏmol^−1^)^15^, yeast (36 ± 3 kJꞏmol^−1^)^16^, and chloroplasts (38 ± 3 and 37 ± 3 kJꞏmol^−1^)^15,16^, supporting the validity of the experiment.

### Cryo-EM structural analysis

We determined the structure of the δ_ΔN_-α fused F_o_F_1_ by single particle cryo-EM analysis. The purified δ_ΔN_-α fused F_o_F_1_ in detergent was applied to EM grids, frozen in liquid ethane, and imaged with 300 kV cryo-EM followed by single-particle analysis using cryoSPARC. The cryo-EM map was obtained by *ab initio* 3D reconstruction and classification followed by refinement with C1 symmetry. As the rotor complex in F_o_F is oriented at one of the three catalytic dwell angles relative to the stator ring and peripheral stalk, when aligned against the central core complex including the rotor complex, the particles showed three positions of the peripheral stalk, each separated by 120° as same as in previous reports^22,23^. Map structure classification was performed by masking the peripheral-stalk positions, confirming the presence or absence of the peripheral stalk at the masked position (Supplementary Fig. 4). This classification was conducted for each stalk position: Stalks 1, 2, and 3. Thus, the map structure was classified into eight sub-classes. The overall resolution was 2.5–3.2 Å (Supplementary Fig. 4). This structural classification confirmed that some fractions of the particles had multiple peripheral stalks (Fig. 5 and Supplementary Fig. 4), two or three of which were not found in the wild-type F_o_F_1_, whereas a significant fraction carried no or a single peripheral stalk. The percentages of F_o_F_1_ structures with 0, 1, 2, and 3 peripheral stalks obtained from the 3D classification were 15, 51, 26, and 8%, respectively. Because of the lower resolution of the F_o_ region, focused refinement with F_o_ was conducted by masking the F_o_ region. The refined structure achieved the resolutions of 3.5–6.6 Å, which confirmed that the peripheral stalk always accompanied F_o_ *a-*subunits. (Supplementary Fig. 4). Thus, the percentage of F_o_F_1_ with 0, 1, 2, and 3 *a*-subunits should correspond to that for peripheral stalks. The fractions of F_o_F_1_ with multiple *a*-subunits were small and the ensemble average of the peripheral stalk is only 1.26/molecule that is evidently lower than the expected value from the subunit stoichiometry analysis and functional analysis of H^+^/ATP ratio, ∼2/molecule. Although the exact reason for this discrepancy is unclear, it is highly likely that the peripheral stalk and the *a*-subunit dissociated due to the meniscus force and/or the interaction with the air/water interface in cryo-EM cells^24^.

**Fig. 5.**
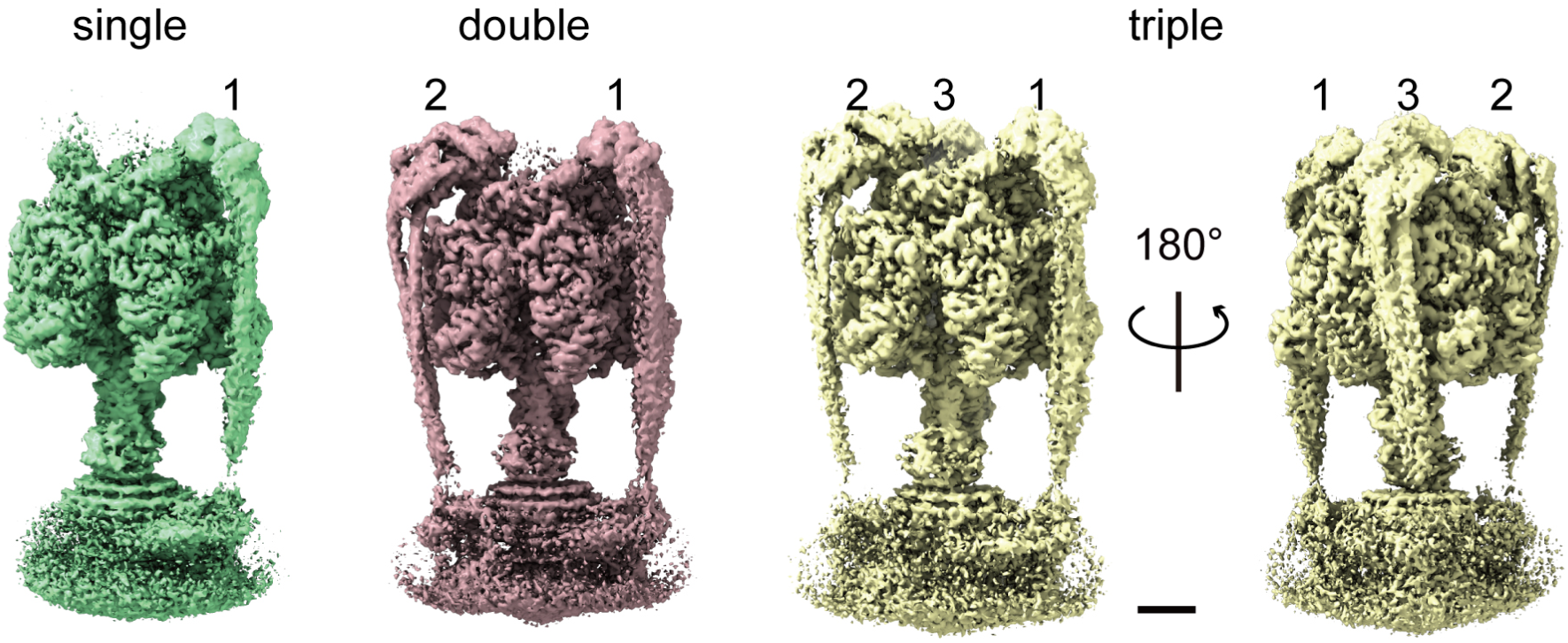
Cryo-EM maps of F_o_F_1_ with multiple peripheral stalks. The cryo-EM maps of F_o_F_1_ with single, double, and triple peripheral stalks. The number represents the position of each peripheral stalk. Scale bar, 25 Å.

### Interaction of peripheral stalk and δ_ΔN_-α fusion

Among the eight sub-classes, there were three structures with a single copy of the peripheral stalk, each at position Stalk1, Stalk 2, or Stalk 3. These structures fit well with the reported three rotational state structure of the wild-type *Bacillus* PS3 F_o_F_1_^22^, respectively (Supplementary Fig. 5), indicating the structural integrity of the binding of the *b*_2_ dimer to F_1_ part via δ_ΔN_-α fusion. The slight differences are found at the top of the F_1_ headpiece and in the side of the central stalk where δ_ΔN_-α fused F_o_F_1_ loses the δ N-terminal domain and the C-terminal helix of the ε subunit. The maps of the double- and triple-stalk F_o_F_1_ were compared with the corresponding maps of single-stalk F_o_F_1_, respectively (Supplementary Fig. 6). These maps were well fitted, indicating that multiple stalks have no significant structural constraints on the whole structure of F_o_F_1_.

The atomic models for triple-stalk F_o_F_1_ are shown in Fig. 6. The structure of δ_ΔN_-α fusion region was well resolved, providing the atomic details on the binding site of δ_ΔN_-α fusion region with the *b*_2_ dimer (Fig. 6a, b). As designed, the three binding sites were almost identical and had basically the same structure found in wild type *Bacillus* PS3 F_o_F_1_^22^ and other ATP synthases^25^ (Supplementary Fig. 7a). These observations reveal that the three *b*_2_ dimers are incorporated via the canonical interaction with δ_ΔN_-*a* fusion, suggesting the integrity of the peripheral stalks of the δ_ΔN_-α fused F_o_F_1_.

**Fig. 6.**
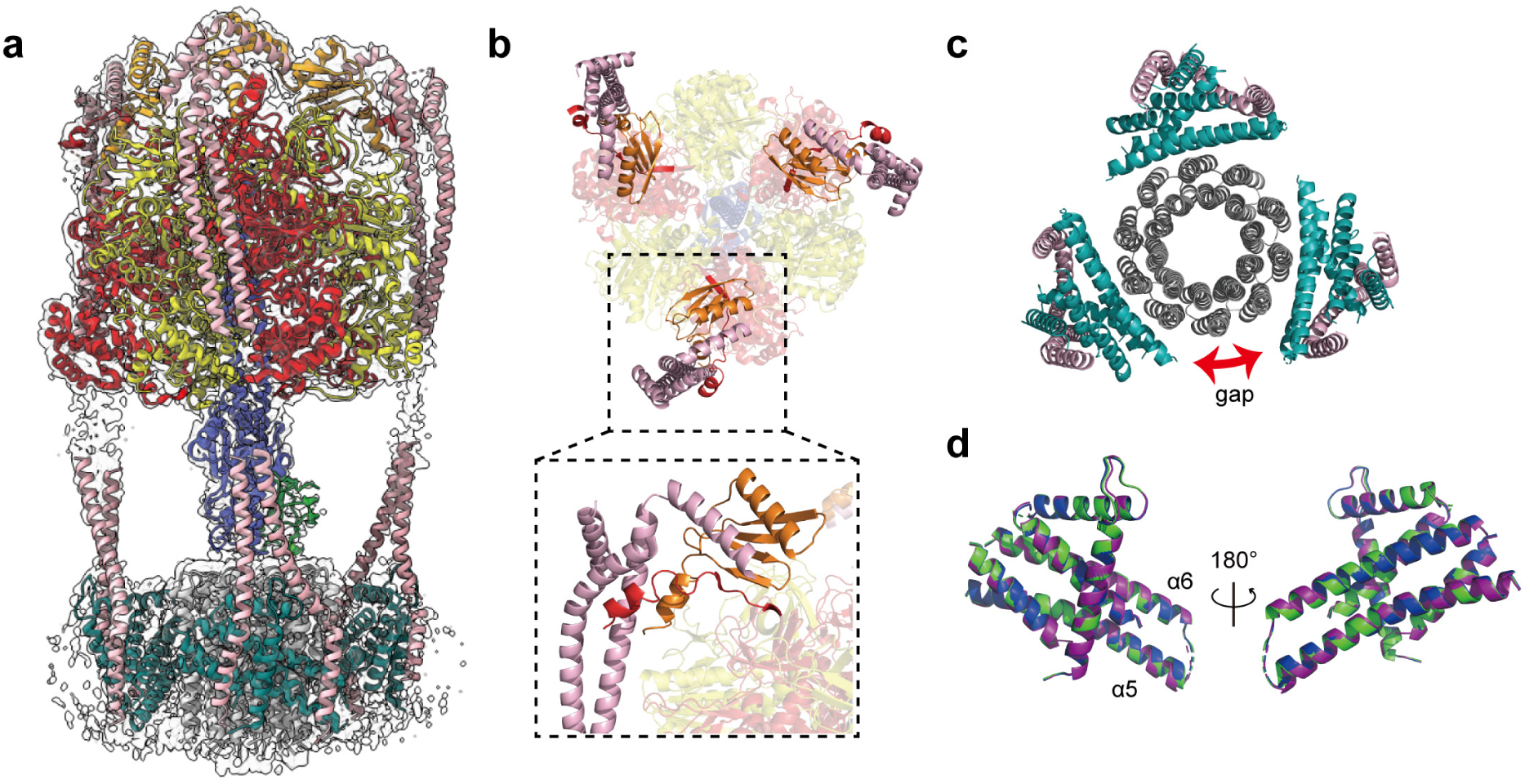
Atomic models of F_o_F_1_ with triple peripheral stalks. (a) Composite map and atomic models for the F_1_ and F_o_ regions of the triple-stalk F_o_F_1_. The middle regions of the *b*-subunits could not be modelled due to the lack of clear density. (b) The top view of the structure. The close-up view shows the side view of the interaction between δ_ΔN_-α fusion and the *b*_2_ dimer. (c) The view from the bottom of the structure shown in (a). (d) The superposition of the three *a-*subunits (green, purple, and blue) in the triple-stalk F_o_.

### Structure of the a-subunits

The structure and the position of the *a*-subunits of the triple-stalk F_o_F_1_ were investigated by comparing them with those of the wild-type *Bacillus* PS3 F_o_F_1_^22^ (Fig. 6c and Supplementary Fig. 7b). The spatial intervals between the *a*-subunits are not perfectly symmetric because of the symmetry mismatch between the ring structures of F_1_ and F_o_, that is, three-fold versus ten-fold. Each *a*-subunit had interactions with the neighboring three *c*-subunits forming an *a*_1_*c*_3_ unit. Therefore, a total of nine *c*-subunits had interactions with the *a*-subunits, leaving the remaining one *c*-subunit at the open position (gap) between two *a-*subunits (Fig. 6c). As a result, the three *a*-subunits were not exactly 120° apart from each other. The asymmetric positioning of the *a*-subunits is in good agreement with that suggested by the three states of wild-type *Bacillus* PS3 F_o_F_1_^22^. When the *a*-subunits in the triple-stalk F_o_F_1_ were compared to each other, Cα-RMSDs were 0.3–0.5 Å (Fig. 6d). In addition, the Cα-RMSDs estimated by superimposition of each *a*-subunit in the triple-stalk F_o_F_1_ with that in the corresponding state of the wild-type F_o_F_1_ were 0.9–1.0 Å. Moreover, superposition of the *a*_1_*c*_3_ units in the same manner gave Cα-RMSDs of 1.0–1.2 Å (Supplementary Fig. 7c). Thus, at the current resolution, the overall structures of the three *a*-subunits in the triple-stalk F_o_F_1_ were principally identical to each other and closely resembled that of the wild-type *Bacillus* PS3 FoF_1_, suggesting that all *a*-subunits are functional, consistent with the abovementioned biochemical analyses showing the enhanced H^+^/ATP ratio.

### Structure of F_1_ part

The structure of the F_1_ portion of the triple-stalk F_o_F_1_ was investigated by comparison with previously reported structures. The F_1_ structure turned out to be very similar to that of *Bacillus* PS3 F_o_F_1_-ε_ΔC_ under uni-site catalysis conditions^23^; one β subunit bound to ADP in a closed conformation 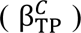 and two β subunits without bound nucleotide taking the open conformations 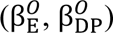 (Supplementary Fig. 8a and b). The purified δ_ΔN_-α fused F_o_F_1_ was prepared in nucleotide-free conditions. Therefore, it is highly likely that bound ADP was endogenous. Notably, at a low-density threshold, weak map density was observed at the outer periphery of 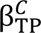, which can be well fitted with the open β conformation of the nucleotide-depleted F_1_ in the *Bacillus* PS3 F_o_F_1_-ε_ΔC_^23^ (Supplementary Fig. 8c). This suggests that the cryo-EM map includes two conformations: the closed and open conformation. The other sub-class structures also exhibited similar mixed maps.

## Discussion

The present study provides significant insights into the design principles of ATP synthases using an engineering approach. Firstly, the δ subunit is the factor that defines the number of peripheral stalks. In this study, the N-terminal domain of δ, which breaks the structural pseudo-threefold symmetry by binding to the position on the symmetry axis, was deleted, and the C-terminal domain of δ was genetically fused to the N-terminal of the α subunit. As a result, up to three peripheral stalks were incorporated into the F_o_F_1_ complex. Structural analysis with cryo-EM revealed that the δ_ΔN_-α fusion and the *b*_2_ dimer took the canonical binding structure as observed in the wild-type F_o_F_1_, except for the absence of the N-terminal domain of the δ subunit. This observation clearly shows that the N-terminal domain of the δ subunit defines the number of peripheral stalks per F_o_F_1_ as one, by breaking the structural symmetry.

Next, the number of the *a-*subunits is determined based on the number of peripheral stalks. While cryo-EM analysis showed that some molecules lost peripheral stalks detaching the *b*_2_ dimer, the observed peripheral stalks always bound to the *a-*subunits, indicating the stable binding between the *a*-subunit and *b*_2_ dimer. Thus, the number of peripheral stalks is the determining factor of the number of the *a*-subunits. In the present study, we did not attempt to introduce more than four peripheral stalks. The structure of the triple-stalk F_o_F_1_ evidently showed that the *c*_10_ ring can accommodate up to three *a*-subunits but not more than four. Considering that each *a-*subunit can interact with three *c*-subunits, it would be possible to accommodate four *a*-subunits in F_o_F_1_ with the *c*-ring composed of more than 12 *c*-subunits.

Another important finding of this study is the functional independence of the *a*-subunit, at least in terms of the coupling stoichiometry of H^+^ (see below). In the triple-stalk F_o_F_1_, all three *a*-subunits interacted with the *c*-ring. In addition, the structural features of the interaction agreed well with those observed in the wild-type F_o_F_1_ structures. This suggests that each of the three *a*-subunits is functional. SDS-PAGE analysis revealed that the samples used in this study had an average of two peripheral stalks and two *a*-subunits. Consistent with these results, analysis of the equilibrium *pmf* showed the doubled H^+^/ATP ratio in comparison with that of the wild type. These results indicate that the coupling stoichiometry of the H^+^ ions is proportional to the number of the *a*-subunits. The additivity in H^+^ stoichiometry means that each *a*-subunit tightly couples H^+^ translocation and rotation of the *c*-ring, regardless of the presence of other *a*-subunits. This is well consistent with the half-channel mode; the model assumes that the probability of H^+^ translocation between the *a*- and the *c*-subunits depends primarily on the relative position of these subunits, which explains well the functional independency of the *a*-subunits. Based on these considerations, we propose that the number of H^+^ ions transported coupled with rotation is determined not only by the number of *c*-subunits constituting the *c*-ring but also by the number of *a*-subunits as follows:

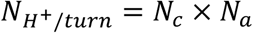

where *N*_H^+^/turn_, *N*_e_, and *N*_a_ represent the total number of H^+^ ions per turn, number of *c-* subunit in the *c-*ring, and number of *a-*subunit in F_o_, respectively. However, one may point to the inconsistency between the biochemical results and the structural analysis with cryo-EM. The proportion of molecules with three peripheral stalks was extremely low, clearly less than the average number indicated by the biochemical results. This is attributable to the dissociation of the peripheral stalks during sample preparation for cryo-electron microscopy. To confirm this point, it is necessary to develop multi-stalk F_o_F_1_ in which peripheral stalks stably bind to F_1_ and to more accurately analyze the correlation between the number of the *a-*subunit and the H^+^ stoichiometry.

As mentioned above, the increased H^+^/ATP ratio suggests the functional independence of the *a-*subunits, but does the independence of the *a*-subunit reaction hold from a kinetic point of view? This raises the question of whether each *a-*subunit proceeds with the H^+^ translocation reaction without mutual interference. Simple reasoning based on the half-channel model suggests that for the *c-*ring to undergo a 36° rotation step, the H^+^ transfer reaction between all *a-c* pairs must be completed. Therefore, the probability of the reaction being completed per unit of time is expected to decrease exponentially as the number of the *a-*subunits increases. Consequently, under conditions where H^+^ transfer between *a-c* pairs is rate-limiting, the rotation speed is expected to decrease with the number of *a-*subunits increases. Indeed, ATP hydrolysis activity and proton pump activity were slower than those of the wild type, suggesting that interference occurs among the *a-*subunits (Supplementary Table 1). However, biochemical experiments are affected by various factors such as the heterogeneity of the sample and the ADP inhibition state, which hamper accurate discussions. Quantitative analysis by single-molecule rotation measurement is necessary to verify this issue.

Based on the findings of this study, we investigated the genes encoding ATP synthases in various species to explore the possibility that ATP synthases with multi-stalk structures exist in nature. We found that the gene operon of ATP synthase from *Acidaminococcus fermentans* shows the gene fusion of the δ and the α subunits (Supplementary Fig. 9). In addition, the N-terminal domain of δ is missing. These features are well consistent with the δ_ΔN_-α fused F_o_F_1_ we designed. Thus, it is highly likely that *A. fermentans* F_o_F_1_ has a multi-stalk structure and can synthesize ATP under low *pmf* conditions. We also found that other species show similar features (UniProt ID: G4Q3K6, A0A1I2C5T3), suggesting more possibility of a multi-stalk F_o_F_1_ in nature. Thus, the present study demonstrates the unexpected diversity of design principle of the F_o_F_1_ ATP synthase which awaits experimental verification.

## Data availability

Cryo-EM maps and models generated in this study were deposited to EMDB (EMD-XXXX to XXXX) and PDB (XXXX and XXXX).

## Acknowledgments

We thank all the members of our laboratory for their helpful comments. We also thank M. Kawasaki, A. Ikeda, S. Inaba, T. Moriya and the staff at the KEK Structural Biology Research Center for their assistance in collecting and analyzing cryo-EM data. We also thank T. Matsui for technical assistance with the biochemical experiments. This study was supported in part by a Grant-in-Aid for Scientific Research on Innovation Areas (JP21H00388 to H.U.), and a Grant-in-Aid for Challenging Research (Exploratory; JP23K18092 to H.U.), and a Grant-in-Aid for Scientific Research (S) (JP19H05624 to H.N.), and a Research Grant from Human Science Frontier Program (Ref. No: RGP0054/2020 to H.N.), and a Research Support Project for Life Science and Drug Discovery (Basis for Supporting Innovative Drug Discovery and Life Science Research (BINDS)) from AMED under Grant Number JP23ama121013 (support number 4318) to T.M. and JP21am0101071 (support number 3071) to T.S.

## Author contributions

H.U., K.Y., and R.M. performed the biochemical experiments. N.H-S. and N.A. performed the cryo-EM analysis. T.S. and T.M. provided the technical support and conceptual advice. H.U. and H.N. conceived and supervised the study and wrote the manuscript. All the authors discussed the results and commented on the manuscript.

## Methods

### Preparation of F_o_F_1_

In this study, *Bacillus* PS3 F_o_F_1_-ε_ΔC_ which has the 10× His-tag at the N-terminus of the β subunit and lacks the inhibitory C-terminal domain of the ε subunit^14,21^ was used as wild-type. The engineered δ_ΔN_-α fused construct of *Bacillus* PS3 F_o_F_1_-ε_ΔC_ lacks the full length δ subunit, and has the δ_ΔN_-α fused subunit in which the N-terminal domain (residues 2-104) of the δ subunit is deleted and its C-terminus is fused to the N-terminus (without Met) of the α subunit with a short linker (GSGG). The wild-type and engineered F_o_F_1_s were expressed in *E. coli* DK8 cells, which lack endogenous F_o_F_1_ genes, by incubating in Super broth at 37℃ for 20 h. Cultured cells were suspended in a solution (10 mM HEPES, pH 7.5, 5 mM MgCl_2_, 10% (v/v) glycerol) and disrupted by sonication. After removing the cell debris at 9,100×g for 45 min, membrane fraction was collected by centrifugation for 131,500×g for 1 h at 4℃. F_o_F_1_ was solubilized from the membrane fraction by the addition of 0.5% (w/v) LMNG (NG310; Anatrace, USA) and incubating for 30 min at 30℃. After centrifugation at 162,000×g for 30 min, the solubilized fraction was applied to a Ni-Sepharose column pre-equilibrated with M buffer (20 mM potassium phosphate buffer and 100 mM KCl, pH 7.5) containing 0.005% LMNG. The column was washed with M buffer containing 20 mM imidazole and 0.005% LMNG, and F_o_F_1_ was eluted with M buffer containing 200 mM imidazole and 0.005% LMNG. The eluted F_o_F_1_ fractions were concentrated before being applied to a Superdex 200 Increase 10/300 column (Cytiva) equilibrated with gel filtration buffer (20 mM HEPES, pH7.5, 100 mM NaCl, and 0.005% LMNG). The peak fractions corresponding to F_o_F_1_ were collected and concentrated to 5–10 mg/mL, frozen with liquid nitrogen, and stored at −80 °C until use. The protein concentrations were determined using a BCA protein assay kit (Pierce) with bovine serum albumin as a standard. The molecular weight of the protein was calculated based on the sequence and subunit stoichiometry. For the δ_ΔN_-α fused F_o_F_1_, the molecular weight was calculated assuming that this F_o_F_1_ had an average of two peripheral stalks.

### Measurement of ATP synthesis/hydrolysis activity of F_o_F_1_

ATP synthesis/hydrolysis activity of F_o_F_1_ was measured using a luciferin-luciferase system at 25°C, as described previously^14^. F_o_F_1_-reconstituted proteoliposomes (PLs) were prepared as described^14^. Then, 300 µL of the proteoliposomes (PLs) were mixed with the 700 µL of acidic buffer containing 50 mM MES or HEPES buffer, 0.143–14.3 mM NaH_2_PO_4_, 6.7 mM KCl, 49 mM NaCl, 4 mM MgCl_2_, 600 mM sucrose and NaOH to obtain the desired pH, and then ADP and valinomycin were added to a final concentration of 20-640 µM and 200 nM, respectively. After incubation for 10–24 h at 25 °C for acidification, base assay medium was prepared by mixing 25 µL of the luciferin/luciferase mixture (2× concentration of CLSII solution in ATP bioluminescence assay kit, Roche, and 5 mM luciferin), 800 µL of the base buffer (380 mM HEPES buffer, 0.1125–11.25 mM NaH_2_PO_4_, 5.63 mM KCl, 55 mM NaCl, 4 mM MgCl_2_, KOH to adjust K^+^ concentration and NaOH to adjust pH), 50-100 µL of ATP and ADP to obtain the desired concentration, and water to adjust total volume to 900 µL and incubated for 10 min for equilibrium. Then the 100 µL of acidified PLs was injected into the base assay medium to initiate the ATP synthesis reaction and the ATP synthesis/hydrolysis activity was monitored with the luciferin/luciferase assay system using a luminometer (Luminescencer AB2200, ATTO). For calibrating luminescence light intensity to ATP concentration, 10 μL of 10 μM ATP was added. The rate was determined from the initial slope of the linear regression of the time courses. The ΔpH was obtained by subtracting pH_in_ from pH_out_, which was determined by directly measuring the pH using a glass electrode. Transmembrane electrical potential, Δψ, was estimated from the Nernst equation.

### Other assays

ATPase activity measurements of PLs using an ATP regeneration system were performed at 25°C in the ATPase assay solution (50 mM HEPES-KOH, pH 7.5, 100 mM KCl, 5 mM MgCl_2_, 2 mM ATP, 1 μg/ml Carbonyl cyanide-p-trifluoromethoxyphenylhydrazone (FCCP), 2.5 mM phosphoenolpyruvate, 100 µg/mL lactate dehydrogenase, 100 µg/mL pyruvate kinase and 0.2 mM NADH) as described previously^26^. ATP-driven H^+^-pumping activity was measured by quenching of ACMA (9-amino-6-chloro-2-methoxyacridine) fluorescence at 25°C in PA4 buffer (10 mM HEPES-KOH, pH 7.5, 100 mM KCl, 5 mM MgCl_2_) supplemented with 0.3 μg/ml ACMA and 1.0 μg/ml F_o_F_1_-reconstituted PLs^26^.

### Cryo-EM grid preparation and data collection

After adding 0.05% lysophosphatidylcholine (1-palmitoyl-2-hydroxy-sn-glycero-3-phosphocholine), 3.0 µL of purified protein (3-6 mg/mL) was loaded onto the glow-discharged Quantifoil R1.2/1.3 grids using a Vitrobot Mark IV (Thermo Fisher Scientific). Grids were blotted for 5 s with a blotting force of 15 under 100% humidity at 18℃, and flash-frozen in liquid ethane. Data were collected using a 300 kV Titan Krios electron microscope (Thermo Fisher Scientific) with a Falcon 4i direct detector device camera with Selectris-X automated with EPU software. Images were recorded in electron counting mode by recording 50 movie frames with an exposure rate of 1.0 e^-^/Å^2^ per frame. The defocus range was 0.8–2.0 μm, and the original pixel size was 0.75 Å.

### Cryo-EM data processing

All image processing steps were performed using cryoSPARC. Details of the image processing workflow are described in Supplementary Fig. 4. A total of 56,081 micrographs were first motion corrected and the CTF was estimated by patch CTF estimation. Particles were manually picked, and templates for particle selection were generated from 2D classification. After template picking, selected 4,316,627 particles were subjected to 2D classification. Then, further selections with ab initio 3D reconstruction, heterogeneous, homogeneous, and non-uniform refinements were performed. All particles after non-uniform refinement were sequentially subjected to a focused 3D refinement with a mask of each of three peripheral stalks including the N-terminal region of the α subunit, C-terminal region of the δ subunit and the hydrophilic region of the *b*_2_-subunits. Each mask was generated from three rotational states of wild-type *Bacillus* PS3 F_o_F_1_ (PDBs 6N2Z, 6N30, and 6N2Y). Eight classes, including one to three peripheral stalks and three rotational states of F_o_F_1_, were identified. The three datasets were collected, merged and refined with non-uniform refinement, resulting in an overall resolution of 2.5–3.2 Å. Further refinement with the F_o_ mask resulted in a map of the F_o_ region with an improved resolution of 3.5–6.6 Å. The resolution was estimated using the FSC criterion of 0.143 threshold. The cryo-EM data collection and refinement statistics were shown in Supplementary Table 2 and 3. The FSC curves and orientation distribution plots were shown in Supplementary Fig. 10 and 11.

### Model building

Models were built and refined in COOT and PHENIX using PDB 6N2Z, 6N30 and 6N2Y as the initial model. Validation statistics were shown in Supplementary Table 2 and 3. Composite map was generated by combining the F_1_ region of the triple-stalk F_o_F_1_ with the F_o_ region of the map from local refinement with F_o_ mask using UCSF Chimera for illustration purpose only. Figures were prepared using the PyMOL, UCSF Chimera, and UCSF ChimeraX. RMSD values for Cα-atoms were calculated using PyMOL align command without outlier rejection.

## SUPPLEMENTARY INFORMATION

**Supplementary Fig. 1.**
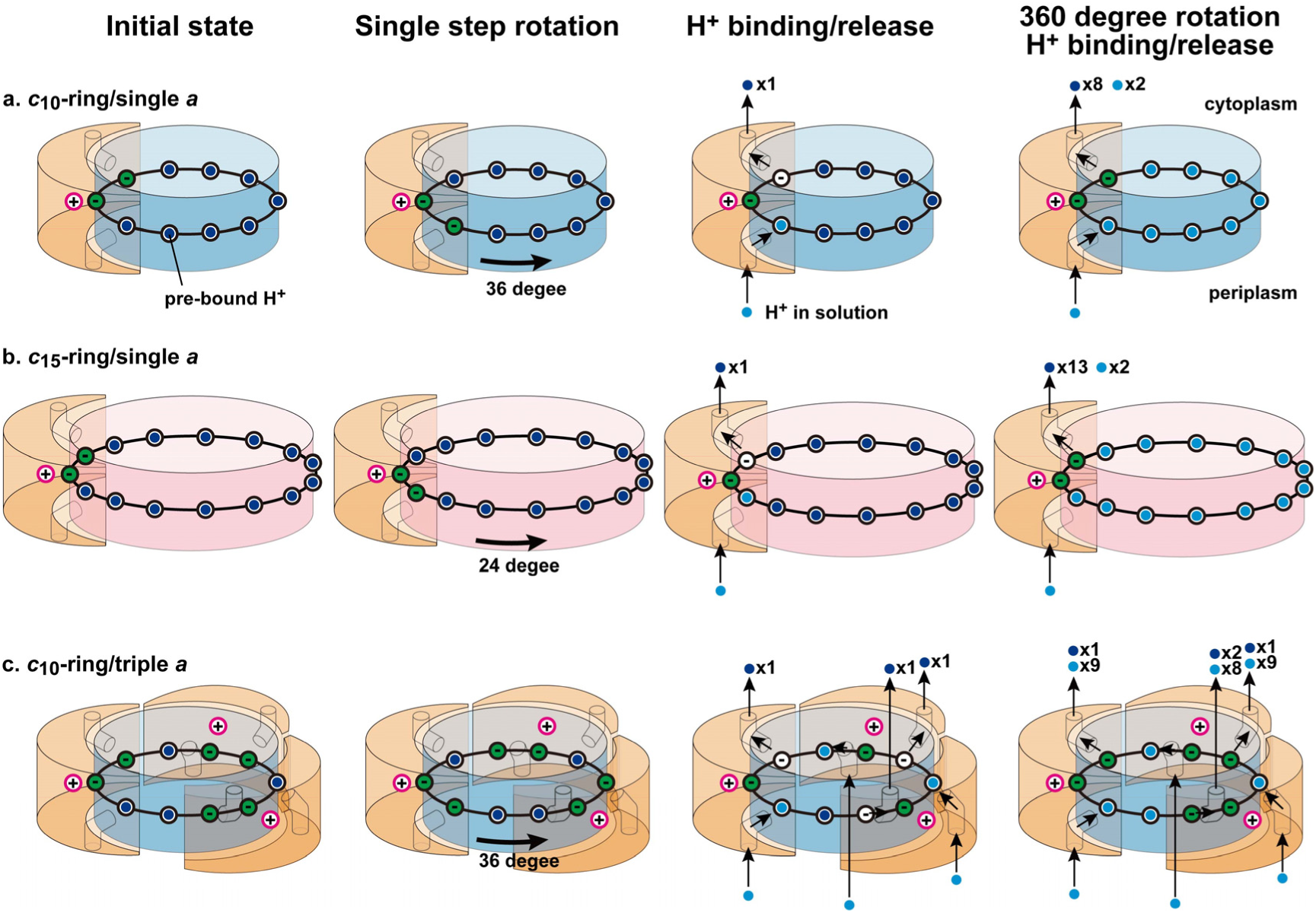
Models of H^+^ translocation through F_o_ per one revolution. The highly conserved arginine residues of *a*-subunit (orange) are depicted with pink open circles. The *c*-subunits are depicted as black open circles with pre-bound H^+^ at the initial state (deep blue circles) and H^+^ incorporated from solution (light blue circles) and, for clarify, deprotonated *c*-subunits facing the *a*-subunit at the initial state are depicted as green circles. (a) Model with *c*_10_-ring and single *a*-subunit. At the initial state, two *c*-subunits are deprotonated. After single step rotation, pre-bound H^+^ is released into the cytoplasmic half-channel due to its interaction with the positively charged arginine residue of *a*-subunit, and the H^+^ in the periplasm enters through the periplasmic half-channel and is transferred to the deprotonated *c*-subunit. In this process, one H^+^ is transferred from the periplasm to the cytoplasm and a total of 10 H^+^s are transferred in one revolution. (b) Model with *c*_15_-ring and a single *a*-subunit. A total of 15 H^+^s are transferred in one revolution. (c) Model with a *c*_10_-ring and triple *a*-subunits. A total of 30 H^+^s are transferred in one revolution.

**Supplementary Fig. 2.**
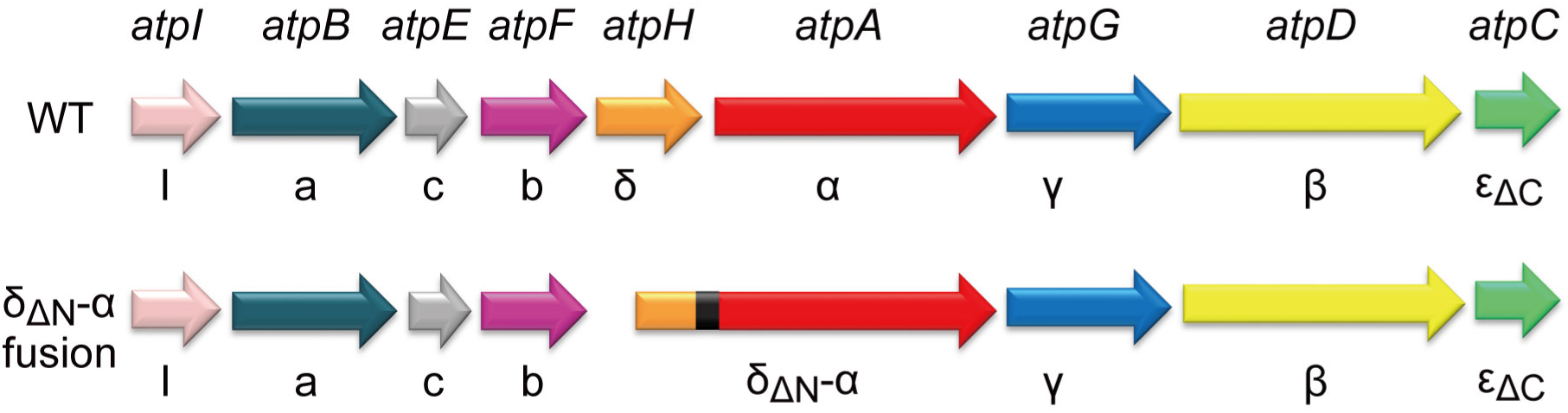
Gene constructs used in this study. *Bacillus* PS3 F_o_F_1_-ε_ΔC_ was used as a wild-type F_o_F_1_. The δ_ΔN_-α fusion mutant lacks the N-terminal domain (residues 2-104) of the δ subunit and the C-terminus was fused to the N-terminus of the α subunit with a short linker sequence (GSGG).

**Supplementary Fig. 3:**
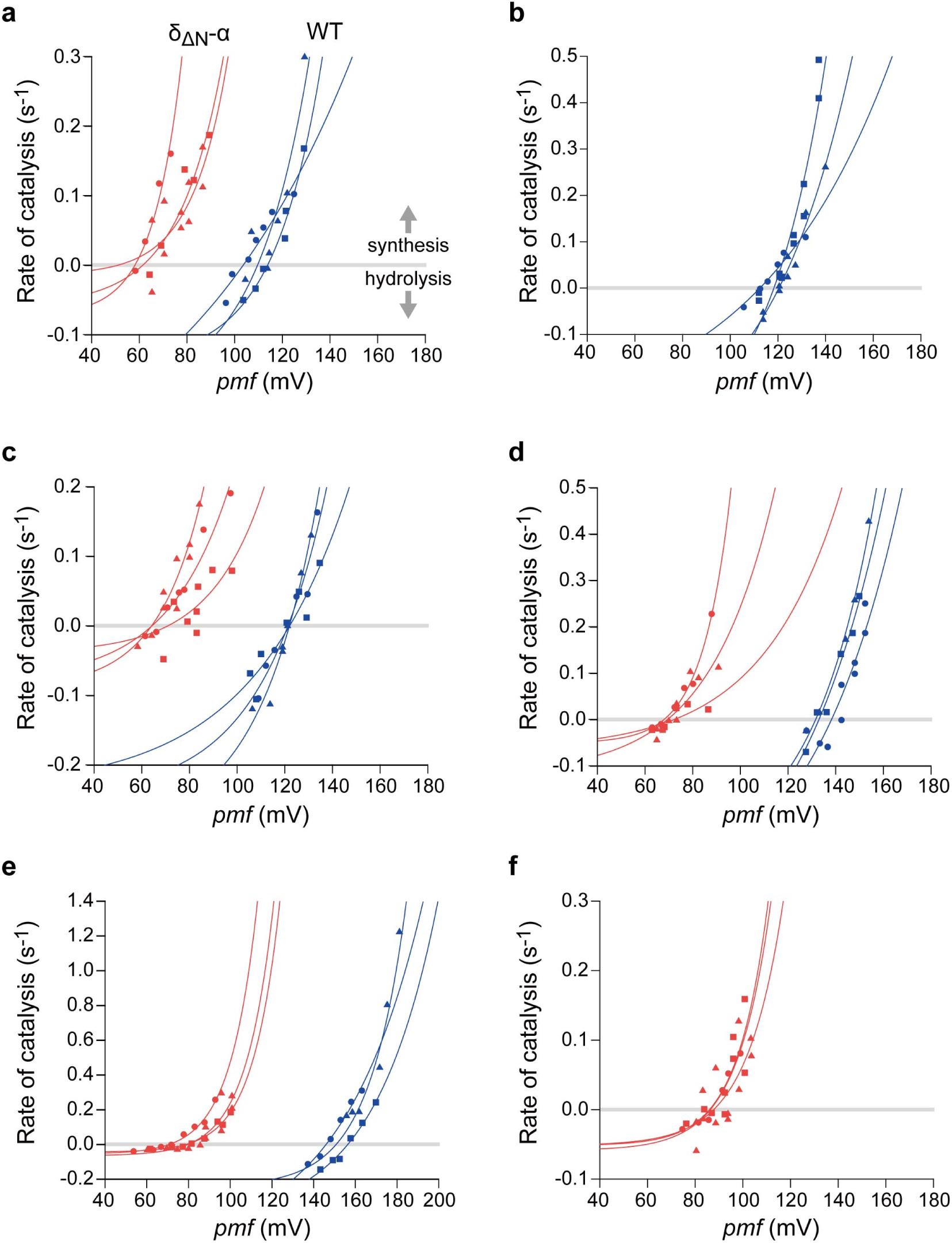
Comparison of the *pmf* dependence of the rate of catalysis at different Q conditions. The results of three repeated experiments (PLs-reconstitutions and measurements) at each *Q* condition are shown. The plots for the δΔN-α fused F_o_F_1_ (red) and wild-type F_o_F_1_ (blue) are shown. (a) *Q* = 0.078; [ATP] = 500 nM, [ADP] = 640 μM, [Pi] = 10 mM. (b) *Q* = 0.208; [ATP] = 500 nM, [ADP] = 240 μM, [Pi] = 10 mM. (c) *Q* = 0.625; [ATP] = 500 nM, [ADP] = 80 μM, [Pi] = 10 mM. (d) *Q* = 2.5; [ATP] = 500 nM, [ADP] = 20 μM, [Pi] = 10 mM. (e) *Q* = 12.5; [ATP] = 500 nM, [ADP] = 40 μM, [Pi] = 1 mM. (f) *Q* = 62.5; [ATP] = 500 nM, [ADP] = 80 μM, [Pi] = 0.1 mM.

**Supplementary Fig. 4.**
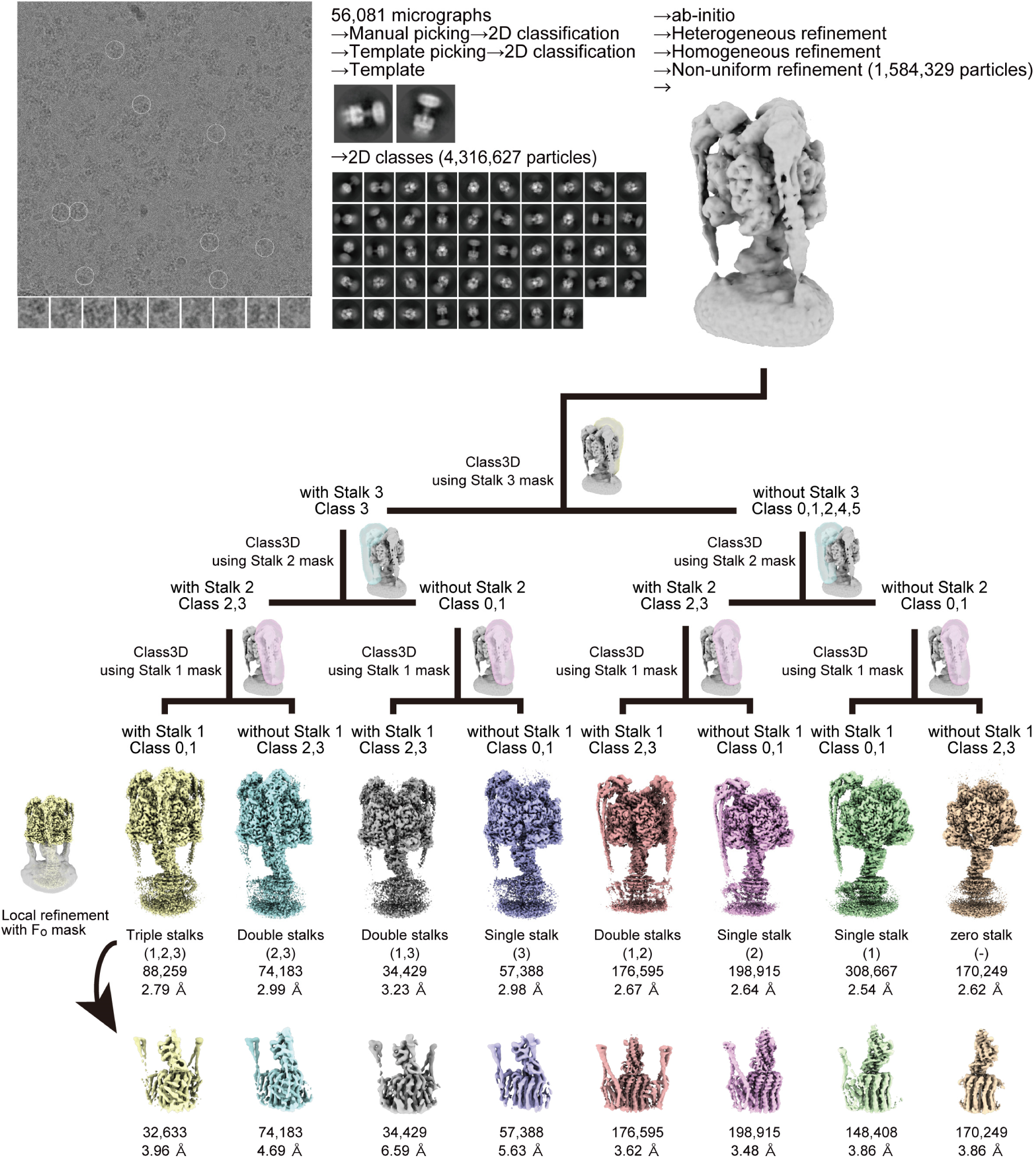
Image processing workflow. Typical micrographs, 2D class averages and the image processing workflow are shown. The masks for Stalk 1, 2, and 3 were generated using the PDB models (6N2Z, 6N30, and 6N2Y), respectively^22^.

**Supplementary Fig. 5.**
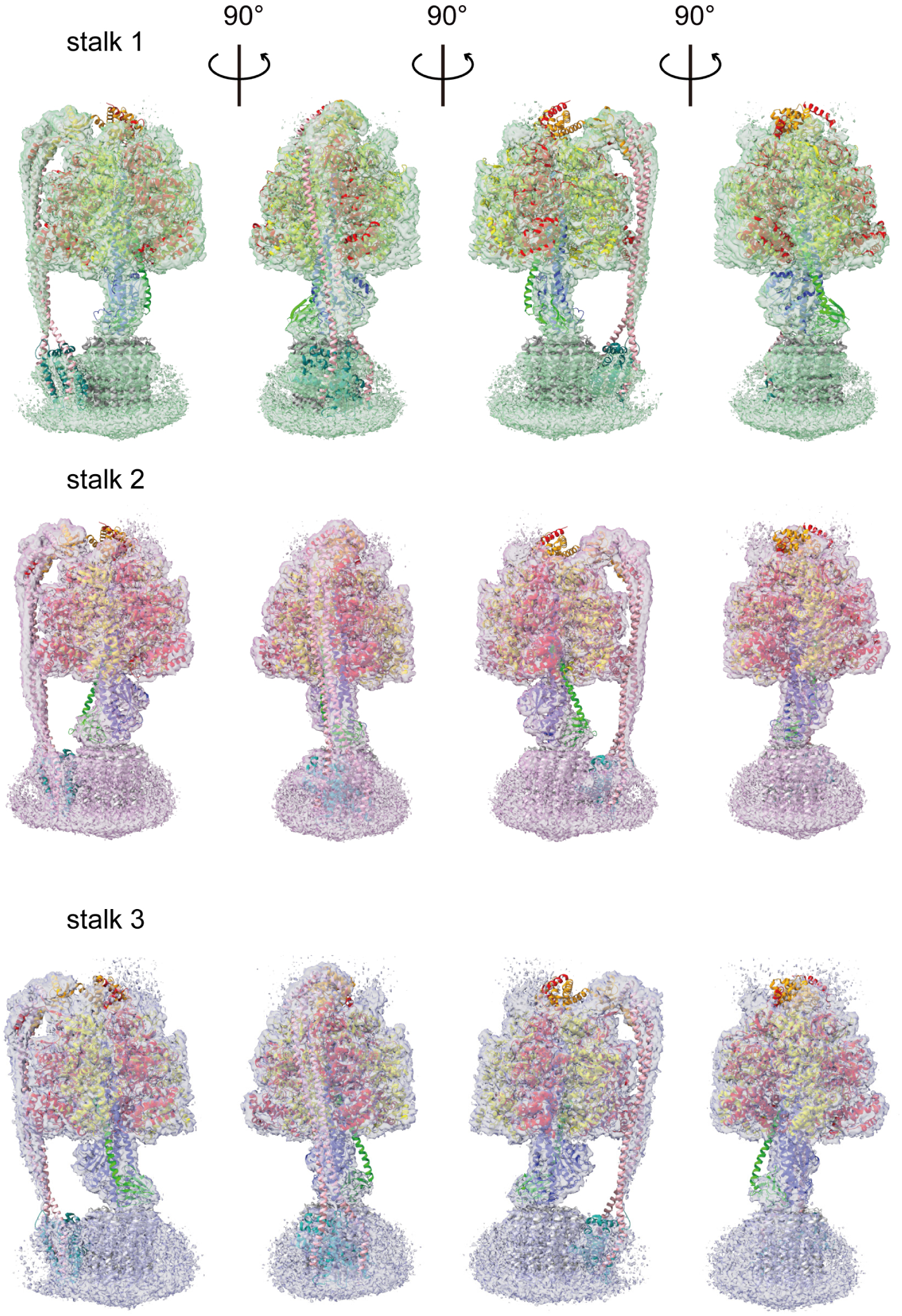
Three rotational states of F_o_F_1_ with single peripheral stalk. The cryo-EM maps of the three single-stalk F_o_F_1_ (Stalk 1, 2, and 3) were fitted with the structures of *Bacillus* PS3 F_o_F_1_ (PDBs 6N2Z, 6N30, and 2N2Y), respectively.

**Supplementary Fig. 6.**
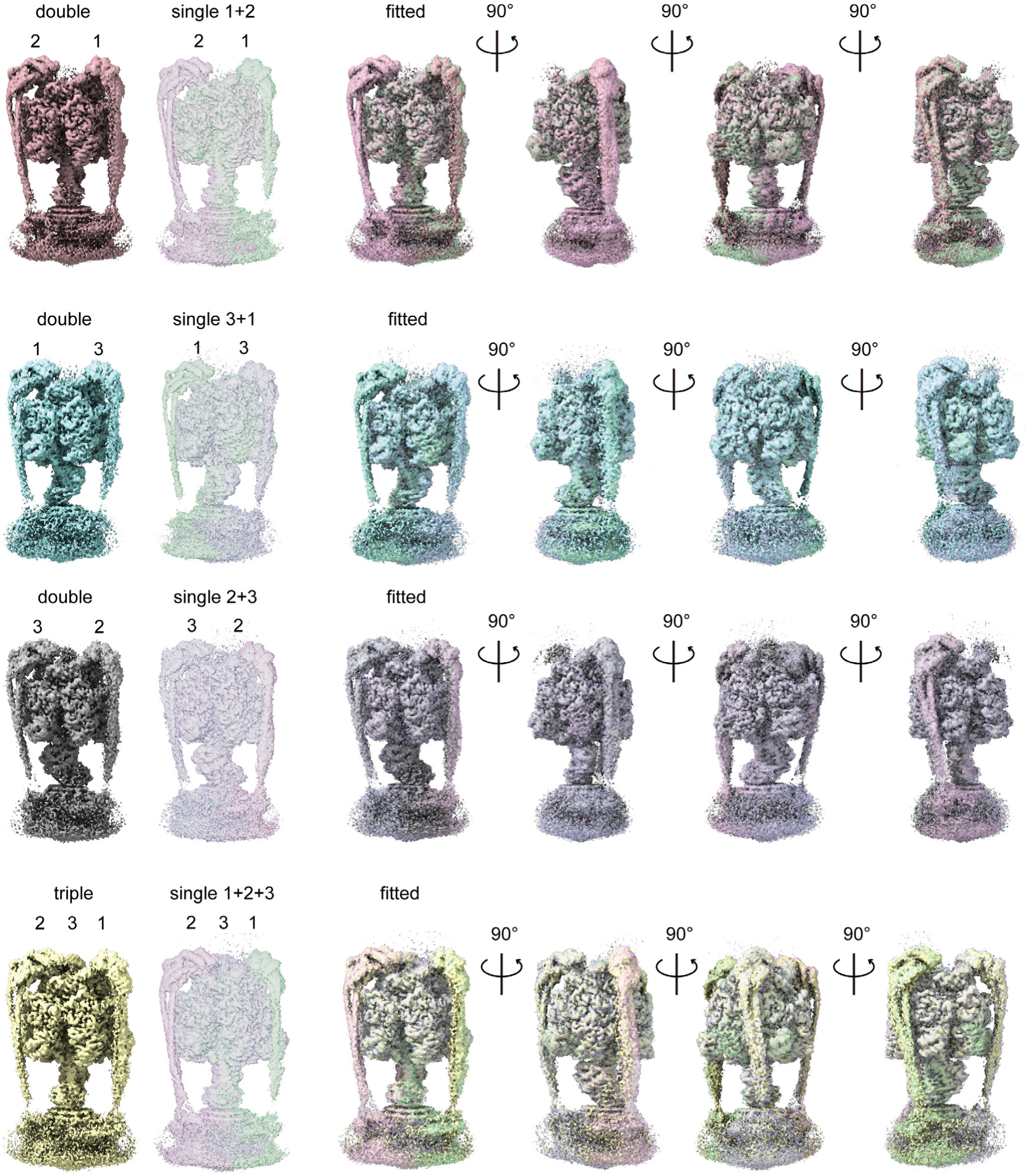
Double- and triple-stalk F_o_F_1_ vs single-stalk F_o_F_1_. The maps of the double- and triple-stalk F_o_F_1_ were fitted with the corresponding maps of single-stalk F_o_F_1_, respectively.

**Supplementary Fig. 7.**
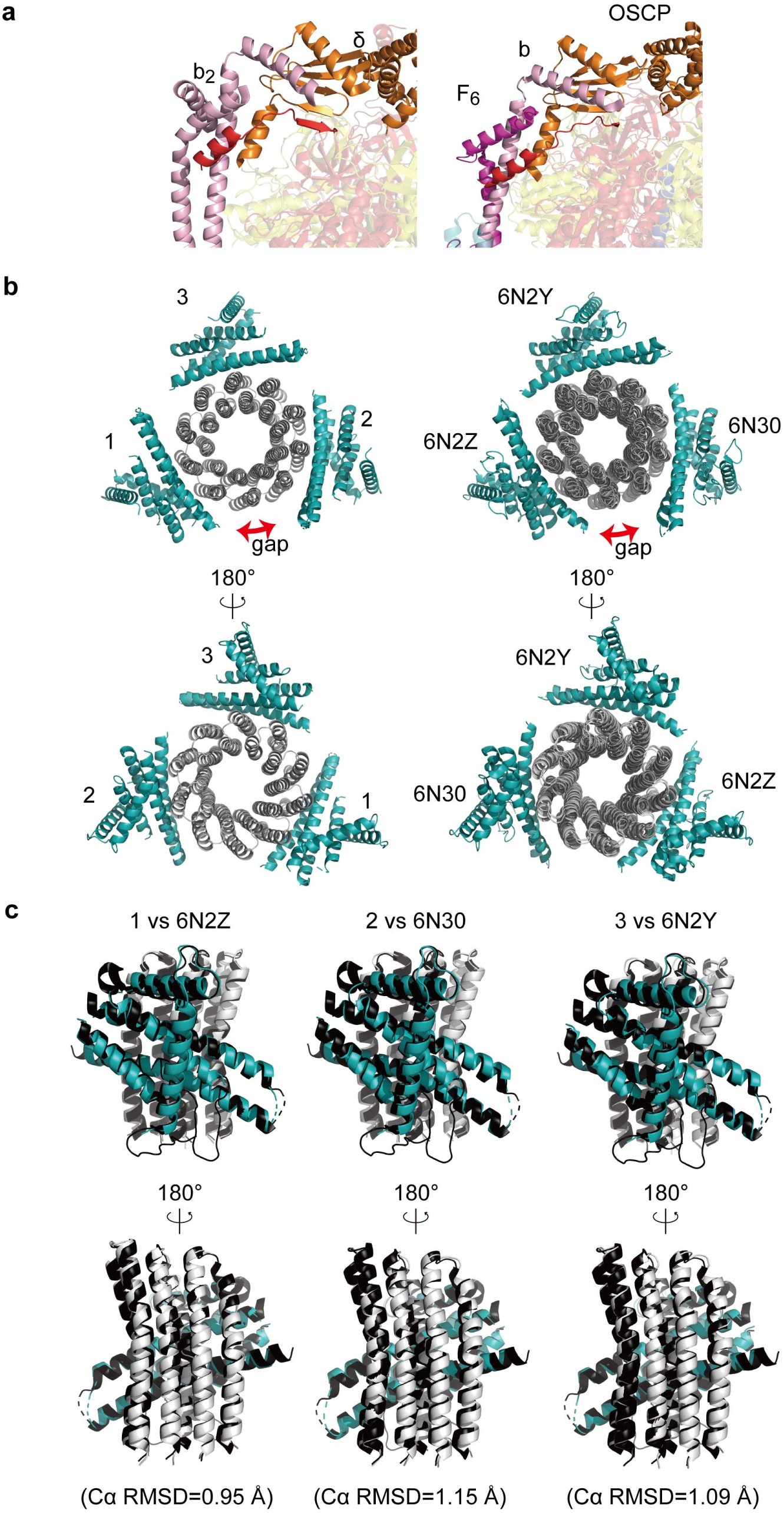
Structural comparison between the triple-stalk F_o_F_1_ and the wild-type F_o_F_1_. (a) The side view of the interaction site between δ (orange), α (red), and *b*_2_ (pink) subunits of the wild-type *Bacillus* PS3 F_o_F_1_ (left, PDB: 6N2Y), and OSCP (orange), α (red), F6 (magenta), and *b* (pink) subunits of bovine F_o_F_1_ (right, PDB: 6YY0). (b) F_o_ of the triple-stalk F_o_F_1_ (left) and the wild-type *Bacillus* PS3 F_o_F_1_ when the three rotational states of the wild-type *Bacillus* PS3 F_o_F_1_ (PDB: 6N2Z, 6N30, and 6N2Y) are aligned with the γ subunit (right). View from the F_o_ side. (c) Superposition of each *a*_1_*c*_3_ unit in the triple-stalk F_o_F_1_ with that in the corresponding state of the wild-type *Bacillus* PS3 F_o_F_1_.

**Supplementary Fig. 8.**
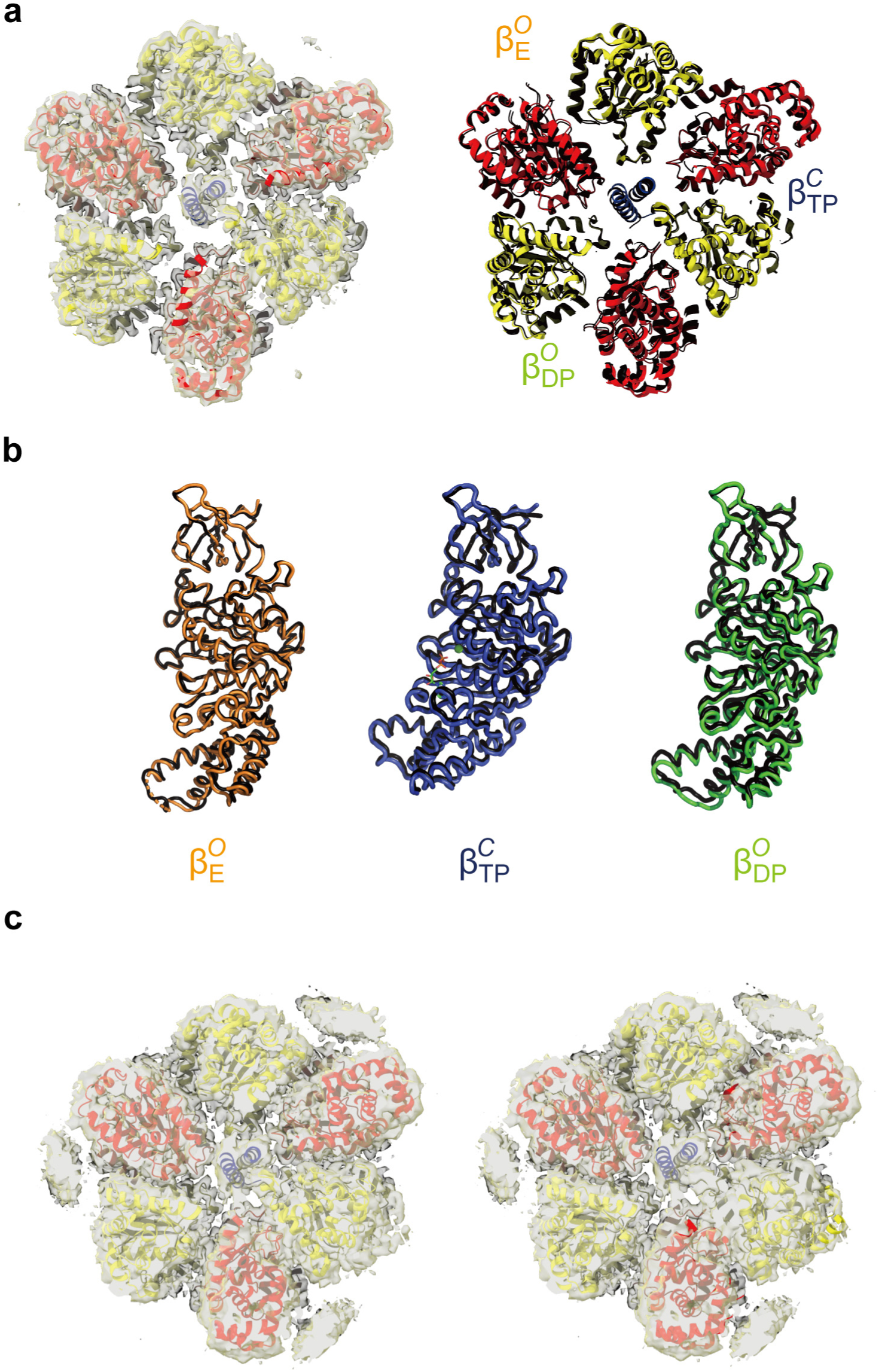
F_1_ part of the triple-stalk F_o_F_1_. (a) The cryo-EM map and the atomic model of the F_1_ part in the triple-stalk F_o_F_1_ viewed from the F_o_ side (left). The α, β, and γ subunits are coloured in red, yellow, and blue, respectively. Superposition of the α_3_β_3_-ring of the triple-stalk F_o_F_1_ (red and yellow) with that of the *Bacillus* PS3 F_o_F_1_-ε_ΔC_ under unisite catalysis conditions (black, PDB: 7XKP) (right). (b) Each β subunit of the triple-stalk F_o_F_1_ 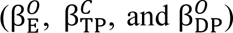 and the *Bacillus* PS3 F_o_F_1_-ε_ΔC_ (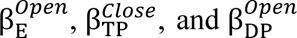, PDB: 7XKP) were superimposed on the N-terminal β-barrel (β2-82). (c) Superposition of the atomic models of the F_1_ part in the triple-stalk F_o_F_1_ (left) and that in the nucleotide-depleted *Bacillus* PS3 F_o_F_1_-ε_ΔC_ (right) onto the cryo-EM map of the triple stalks F_o_F_1_ at lower density threshold than that shown in (a).

**Supplementary Fig. 9.**
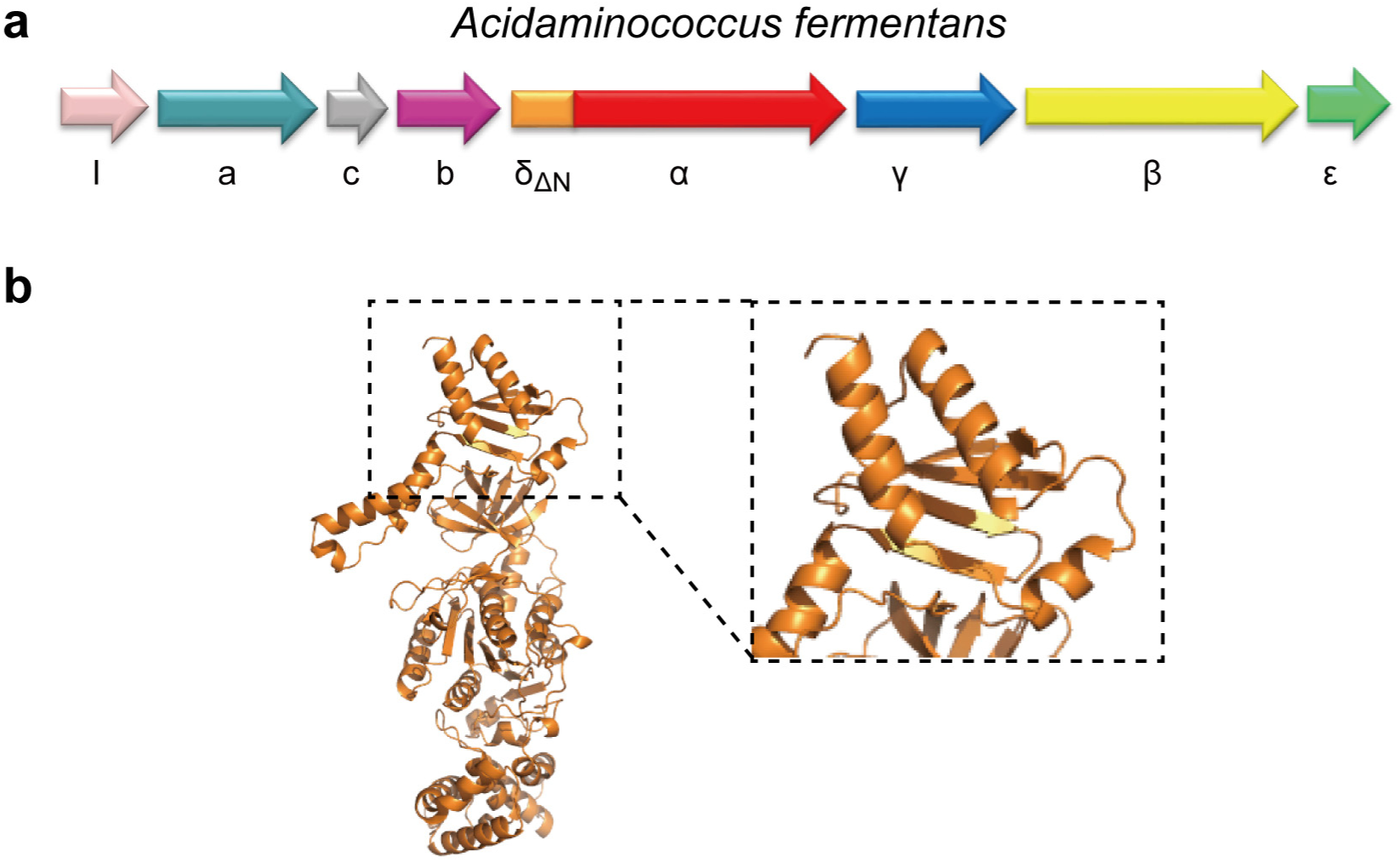
The *atp* operon of *Acidaminococcus fermentans*. (a) The *atp* operon from *Acidaminococcus fermentans* with fused genes for the δ subunit and the α subunit is shown. Notably, the δ asubunit gene lacks the N-terminal domain, similar to the δΔN-α fused F_o_F_1_ developed in the present study. (b) The predicted structures of the δΔN-α subunit of *Acidaminococcus fermentans* from the AlphaFold Protein Structure Database (UniProt ID: D2RLW6). The structure is also similar to the corresponding part of the engineered δΔN-α fused F_o_F_1_.

**Supplementary Fig. 10.**
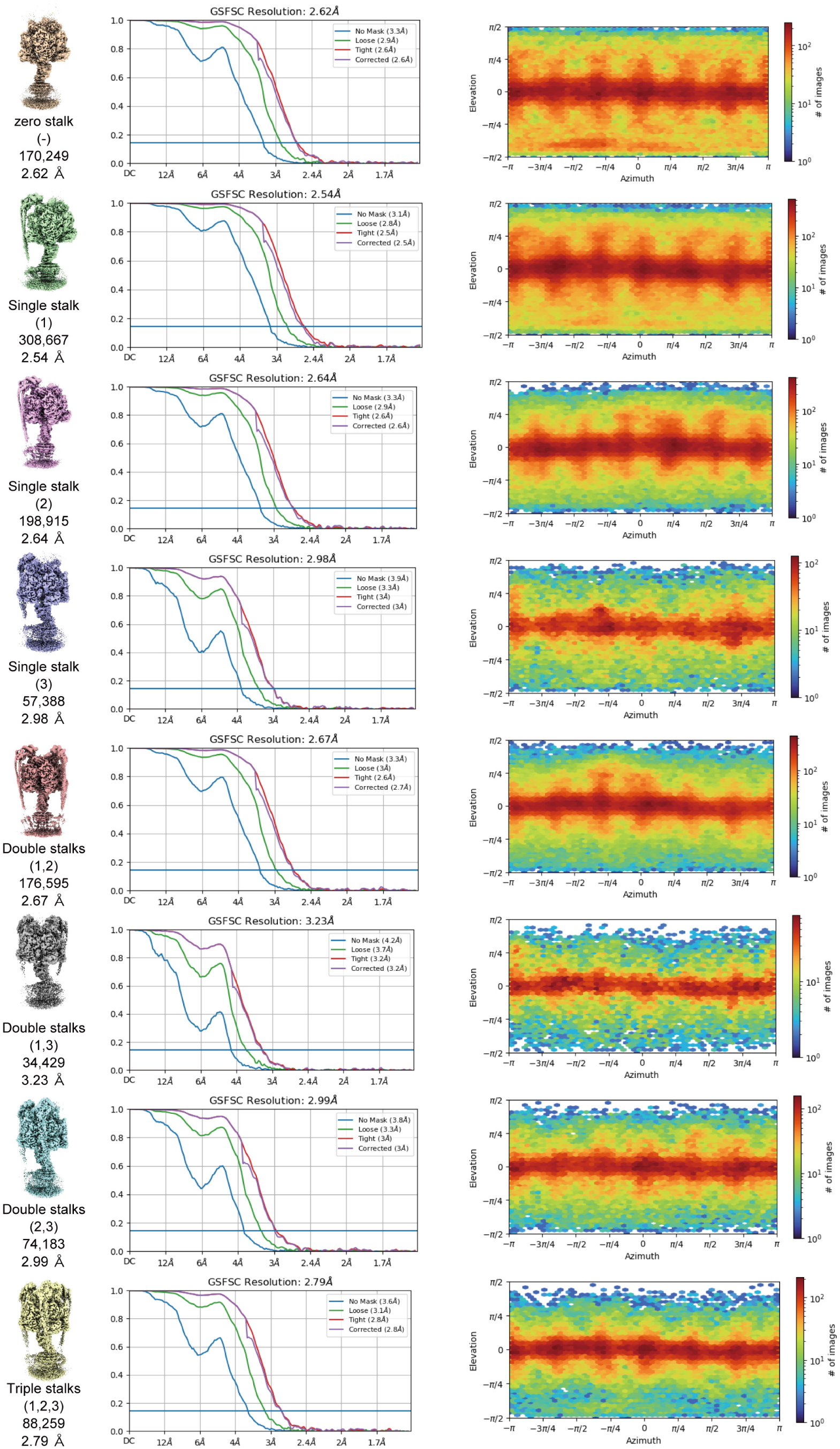
FSC curves. The FSC curves and orientation distribution plots for each F_o_F_1_ map from cryoSPARC.

**Supplementary Fig. 11.**
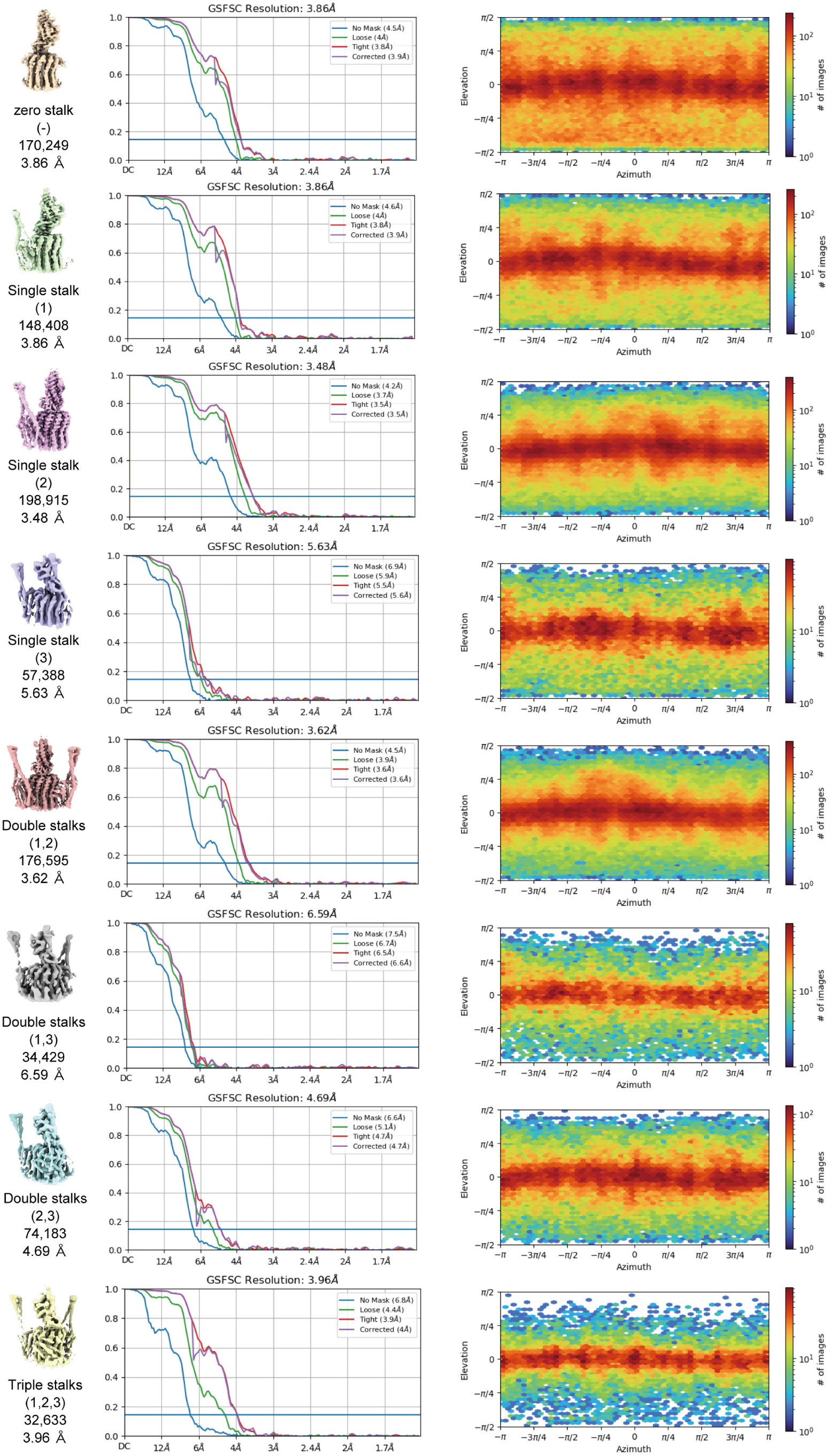
FSC curves. The FSC curves and orientation distribution plots for each F_o_ map from cryoSPARC.

**Supplementary table 1.** H^+^-pumping and ATPase activity of F_o_F_1_ reconstituted proteoliposomes at 2 mM ATP.

| Protein | H<sup>+</sup>-pumping activity | ATPase activity |
| --- | --- | --- |
| Wild-type F<sub>o</sub>F<sub>1</sub> | 75±5% ACMA quench | 53 ± 6 s<sup>-1</sup> |
| δ<sub>ΔN</sub>-α fused F<sub>o</sub>F<sub>1</sub> | 20 ±5% ACMA quench | 21 ± 3 s<sup>-1</sup> |

**Supplementary table 2.**
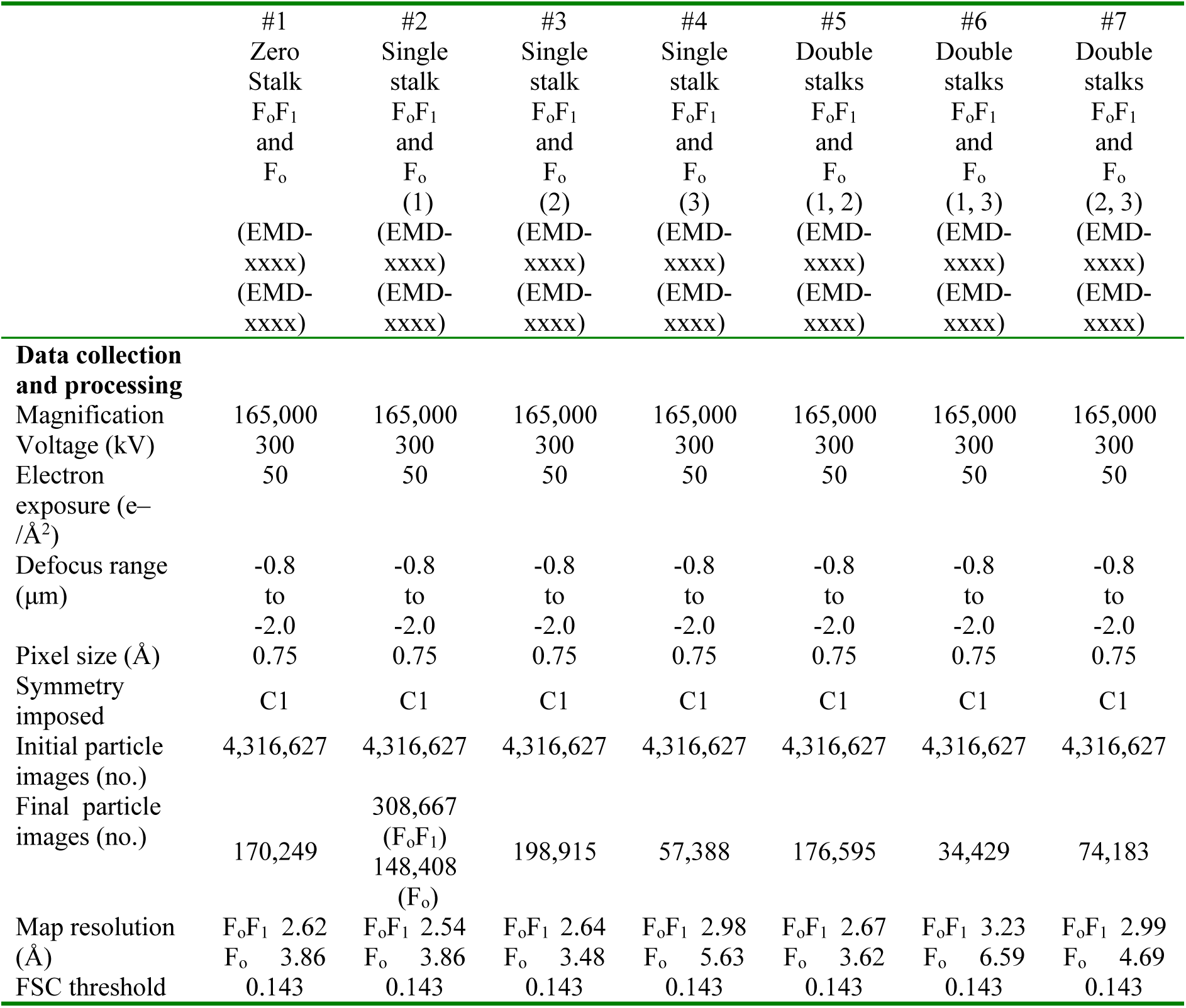
Cryo-EM data collection, refinement and validation statistics for F_o_F_1_ with 0, 1, and 2 peripheral stalks.

**Supplementary table 3.** Cryo-EM data collection, refinement and validation statistics for F_o_F_1_ with triple peripheral stalks.

|  | #8<br>Triple<br>Stalks<br>F<sub>0</sub>F<sub>1</sub><br>(EMD-xxxx) | #9<br>F<sub>1</sub><br>of<br>Triple stalks F<sub>0</sub>F<sub>1</sub><br>(EMD-xxxx)<br>(PDB xxxx) | #10<br>F<sub>0</sub><br>of<br>Triple stalks F<sub>0</sub>F<sub>1</sub><br>(EMD-xxxx)<br>(PDB xxxx) |
| --- | --- | --- | --- |
| **Data collection and processing** |  |  |  |
| Magnification | 165,000 | 165,000 | 165,000 |
| Voltage (kV) | 300 | 300 | 300 |
| Electron exposure (e<sup>-</sup>/Å<sup>2</sup>) | 50 | 50 | 50 |
| Defocus range (μm) | -0.8 to -2.0 | -0.8 to -2.0 | -0.8 to -2.0 |
| Pixel size (Å) | 0.75 | 0.75 | 0.75 |
| Symmetry imposed | C1 | C1 | C1 |
| Initial particle images (no.) | 4,316,627 | 4,316,627 | 4,316,627 |
| Final particle images (no.) | 88,259 | 88,259 | 32,633 |
| Map resolution (Å) | 2.79 | 2.79 | 3.96 |
| FSC threshold | 0.143 | 0.143 | 0.143 |
| **Refinement** |  |  |  |
| Initial model used (PDB code) |  | 6N2Z | 6N2Z |
| Model resolution (Å) |  | 3.1 | 4.3 |
| FSC threshold |  | 0.5 | 0.5 |
| Map sharpening *B* factor (Å<sup>2</sup>) |  | 49.8 | 96.1 |
| Model composition |  |  |  |
| Non-hydrogen atoms |  | 28206 | 10826 |
| Protein residues |  | 3853 | 1875 |
| Ligands |  | 4MG,1ADP | N/A |
| *B* factors (Å<sup>2</sup>) |  |  |  |
| Protein |  | 151.14 | 193.72 |
| Ligand |  | 293.97 | N/A |
| R.m.s. deviations |  |  |  |
| Bond lengths (Å) |  | 0.004 | 0.003 |
| Bond angles (°) |  | 0.569 | 0.611 |
| Validation |  |  |  |
| MolProbity score |  | 1.43 | 1.51 |
| Clashscore |  | 5.47 | 9.79 |
| Poor rotamers (%) |  | 1.02 | 0.56 |
| Ramachandran plot |  |  |  |
| Favored (%) |  | 97.35 | 98.28 |
| Allowed (%) |  | 2.60 | 1.72 |
| Disallowed (%) |  | 0.05 | 0.00 |

